# Ensemblex: an accuracy-weighted ensemble genetic demultiplexing framework for population-scale scRNAseq sample pooling

**DOI:** 10.1101/2024.06.17.599314

**Authors:** Michael R. Fiorini, Saeid Amiri, Allison A. Dilliott, Cristine M. Yde Ohki, Lukasz Smigielski, Susanne Walitza, Edward A. Fon, Edna Grünblatt, Rhalena A. Thomas, Sali M.K. Farhan

## Abstract

Multiplexing samples from distinct individuals prior to sequencing is a promising step toward achieving population-scale single-cell RNA sequencing by reducing the restrictive costs of the technology. Individual genetic demultiplexing tools resolve the donor-of-origin identity of pooled cells using natural genetic variation but present diminished accuracy on highly multiplexed experiments, impeding the analytic potential of the dataset. In response, we introduce Ensemblex: an accuracy-weighted, ensemble genetic demultiplexing framework that integrates four distinct algorithms to identify the most probable subject labels. Using computationally and experimentally pooled samples, we demonstrate Ensemblex’s superior accuracy and illustrate the implications of robust demultiplexing on biological analyses.

## Background

Single-cell RNA sequencing (scRNAseq) continues to revolutionize our molecular understanding of biology by providing unprecedented insight into the transcriptional landscape of individual cells. Unlike bulk RNAseq, where the RNA from all cells within a tissue is sequenced to produce total expressional profiles across all cells, scRNAseq captures transcriptional signatures at a single-cell resolution, elucidating the diverse gene expression across distinct cell types and subtypes. Differential gene expression (DGE) can then be calculated between subgroups of cells to reveal cell type-specific expression changes between patient or treatment groups. However, scRNAseq has come at the expense of increased costs, hindering its application for population-scale analyses, which are critical for deriving clinico-pathological associations and characterizing the genetic heterogeneity of complex diseases in biomedical sciences (1, 2).

In addition to the expense of separately capturing and sequencing cells from individual donors, the costs of scRNAseq are exacerbated for cell cultures, such as those derived from induced pluripotent stem cells (iPSC) (1). In particular, neurological diseases are difficult to study in human tissue because access to post-mortem brains is limited and experimental manipulations are not possible; in contrast, iPSC-derived cultures of neurons and other brain cells grown from reprogrammed skin or blood cells of human donors are an excellent model of the brain (3). However, iPSCs from each donor must be individually plated and differentiated in parallel, presenting prohibitively high consumable and labour costs that render the methodology unfeasible for population-scale analyses. Multiplexing cultures by pooling cells from multiple donors prior to growth and differentiation, droplet capture, and sequencing, is one solution to address this limitation as it reduces costs by a factor of the number of samples multiplexed (4). Similarly, samples such as tumor biopsies can be pooled at acquisition to realize the same benefits. In turn, genetic demultiplexing tools are cost-effective, statistical frameworks that use the natural genetic variation at sites of single-nucleotide polymorphisms (SNP) observed in the transcriptome to cluster cells on the basis of their donor’s genotype. Importantly, genetic demultiplexing can be informed by prior genotype information of the donors to improve demultiplexing accuracy and facilitate the assignment of each cell back to its specific donor-of-origin, which is critical for downstream analyses aiming to investigate discrepancies between subjects. At present, six genetic demultiplexing tools have been developed for scRNAseq: Demuxalot (5) and Demuxlet (6) both require prior genotype information as input; Freemuxlet (6) relies entirely on the de novo transcriptome and does not incorporate prior genotype information; and ScSplit (7), Souporcell (8), and Vireo (9) provide versions of the algorithm that can work with and without prior genotype information **(Table 1).**

**Table 1.**
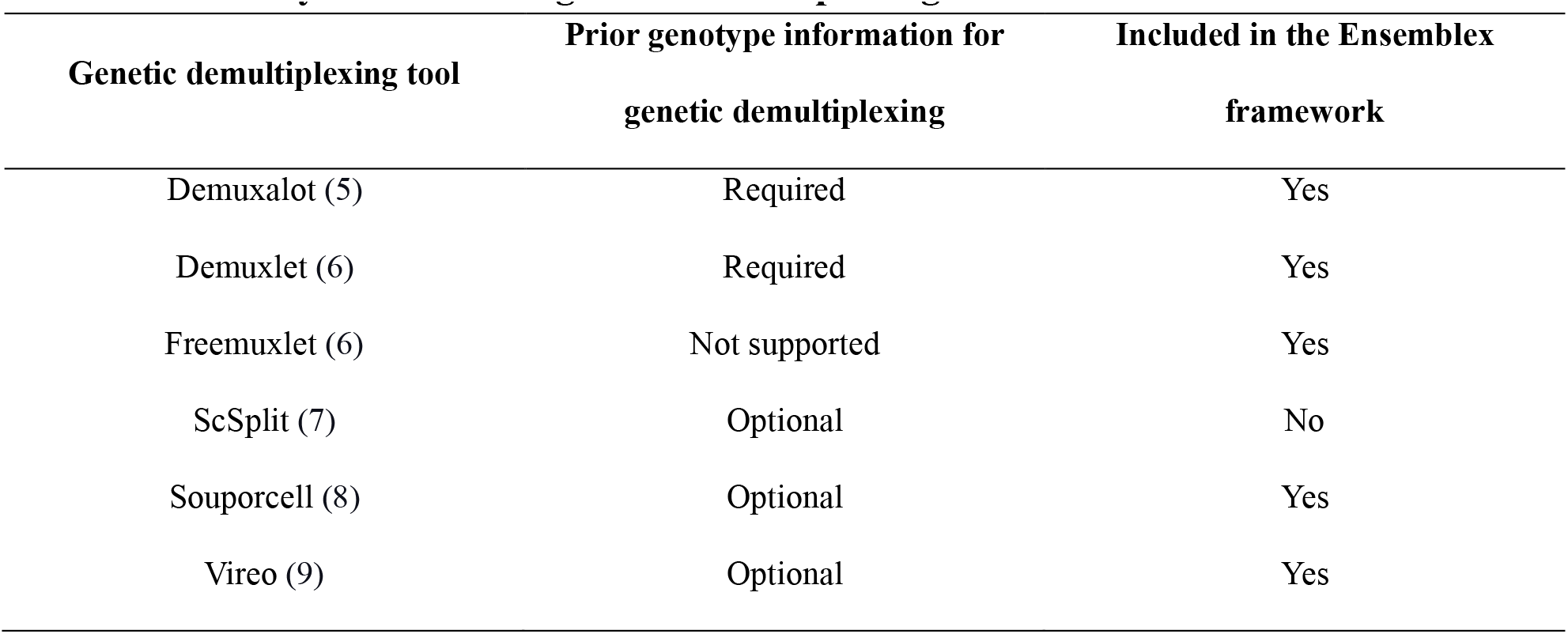
Summary of individual genetic demultiplexing tools.

A robust genetic demultiplexing tool is tasked with mitigating the addition of technical artifacts into scRNAseq datasets by correctly classifying each pooled cell to its donor-of-origin, correctly identifying heterogenic doublets (erroneous barcodes composed of two or more cells from distinct subjects), and quantifying its confidence in the demultiplexed labels so that low-confidence classifications can be eliminated from downstream analyses. While benchmarking analyses on the available genetic demultiplexing tools have shown effectiveness for demultiplexing small sample sizes, limitations emerge as the number of multiplexed samples approach a population scale (6) (7) (8) (9). For example, using computationally multiplexed samples, Neavin et al. evaluated the performance of genetic demultiplexing tools as the number of samples approached a population scale and observed diminished demultiplexing accuracy with increasing numbers of pooled samples, as well as notable classification discrepancies between tools (10). Furthermore, even at small sample sizes, divergent assignments between genetic demultiplexing tools are common (8) (9) (11). Another feature that has been shown to affect genetic demultiplexing performance is the underrepresentation of samples in a pool, which is especially relevant for cell culture-based multiplexed experiments, as variable growth rates *in vitro* across cell lines is common (12) (8) (9). Genetic demultiplexing tools have also shown low concordance for identifying heterogenic doublets, which should be removed prior to downstream analyses to avoid technical noise in the data (10). Importantly, benchmarking analyses have repeatedly highlighted ScSplit’s poor performance relative to the remaining tools (9) (10) (8) (11). The sum of these limitations calls to question the robustness of the individual genetic demultiplexing tools for resolving the donor identities of highly multiplexed samples, which represents an important hurdle for feasibly achieving population-scale scRNAseq analysis.

In response to the divergent assignments commonly observed across tools, a consensus framework, whereby only cells that show matching sample labels across all individual tools are retained for downstream analyses, may appear sufficient to resolve the risk of introducing technical noise into the data from misclassified cells. However, consensus frameworks are restricted to performing only as well as the worst-performing tool, and genetic demultiplexing performance is highly dataset dependent (10); thus, the overall performance of a consensus framework can vary immensely between datasets. To this end, Neavin et al. proposed a majority vote framework for genetic demultiplexing, whereby a cell is assigned to the sample called by the majority of tools (10). However, this approach can be vulnerable to a subset of tools performing poorly on the dataset, does not allocate additional weight to the votes of tools that perform more favourably on the dataset, cannot account for instances when ties occur amongst tools, and cannot capture cells that are correctly classified by only one tool. The sum of these limitations leads to the unnecessary removal of cells from downstream analyses, reducing statistical power, especially for highly multiplexed pools where each donor, on average, will have a lower representation of cells in the pool. Moreover, the ability to capture the transcriptional profiles of rare cell types with scRNAseq provides a notable advancement over bulk RNAseq and can strongly influence biological interpretations (13); thus, investigators are reluctant to discard valuable cells in order to maximize the analytic potential of their dataset.

To address the need for a robust genetic demultiplexing framework that can maximize the number of confidently classified cells retained for downstream analyses, achieve high demultiplexing accuracy for population-scale scRNAseq sample pooling, and maintain reliability across different datasets, we developed Ensemblex: an accuracy-weighted ensemble genetic demultiplexing framework designed to identify the most probable sample labels from each of its constituent tools — Demuxalot, Demuxlet/Freemuxlet, Souporcell, and Vireo. Our ensemble method capitalizes on combining distinct statistical frameworks for genetic demultiplexing while adapting to the overall performance of its constituent tools on the respective dataset, making it resilient against a poorly performing tool and facilitating a higher yield of cells for downstream analyses. The Ensemblex workflow is assembled into a three-step pipeline — 1) accuracy-weighted probabilistic ensemble; 2) graph-based doublet detection; 3) Ensemble-independent doublet detection — and can demultiplex pools with or without prior genotype information.

Here, we showcase Ensemblex’s improved demultiplexing performance across a variety of settings through benchmarking analyses on a total of 141 computationally multiplexed pools with known ground-truth sample labels ranging in size from 4 to 80 samples. We applied the ensemble method to three diverse, experimentally multiplexed datasets: 1) non-small cell lung cancer (NSCLC) dissociated tumor cells from 7 individuals with donor-specific oligonucleotide labels; 2) iPSC-derived dopaminergic neurons (DaN) from 22 healthy individuals; and 3) iPSC-derived neural stem cells (NSC) from 9 individuals with attention deficit hyperactivity disorder (ADHD) and 7 healthy controls. We demonstrate Ensemblex’s robustness across distinct datasets, its ability to return a high proportion of confidently classified cells for downstream analysis, and the implications that its improved demultiplexing performance has on biological interpretations of multiplexed experiments.

## Results and Discussion

### Evaluating the performance of existing individual genetic demultiplexing tools

To evaluate the performance of individual genetic demultiplexing tools, we generated computationally multiplexed pools using scRNAseq of 80 different iPSC lines from Parkinson’s disease patients and healthy controls, which were differentiated towards a DaN state as part of the Foundational Data Initiative for Parkinson’s Disease (FOUNDIN-PD) (14). Processed scRNAseq data from the independent iPSC lines were merged to simulate sample-pooling using a previously described protocol (9), which provided known ground-truth donor and doublet labels. We generated 96 *in silico* pools ranging in size from 4 to 80 multiplexed samples, where each sample corresponded to a unique donor-of-origin. The *in silico* pools averaged 17,396 cells per pool with a constant 15% doublet rate.

Leveraging whole-genome sequencing (WGS) of the 80 donors from which the iPSC lines were derived and the four genetic demultiplexing tools that can utilize prior genotype information — Demuxalot, Demuxlet, Souporcell, and Vireo-GT — we first investigated the proportion of correctly classified cells by the individual tools (**Figure 1A**). Across the 96 *in silico* pools, all tools showed decreasing demultiplexing performance as the number of samples within the pool increased. Souporcell demonstrated the largest decrease in the proportion of correctly classified cells as the number of multiplexed samples increased from 4 (mean = 90.60%) to 80 (mean = 53.27%). In accordance with previous findings (10, 15), the individual genetic demultiplexing tools performed better on singlet classification than doublet detection, highlighting an avenue for improved genetic demultiplexing accuracy by increasing the rate of heterogenic doublet identification (**Figure 1A**).

**Figure 1.**
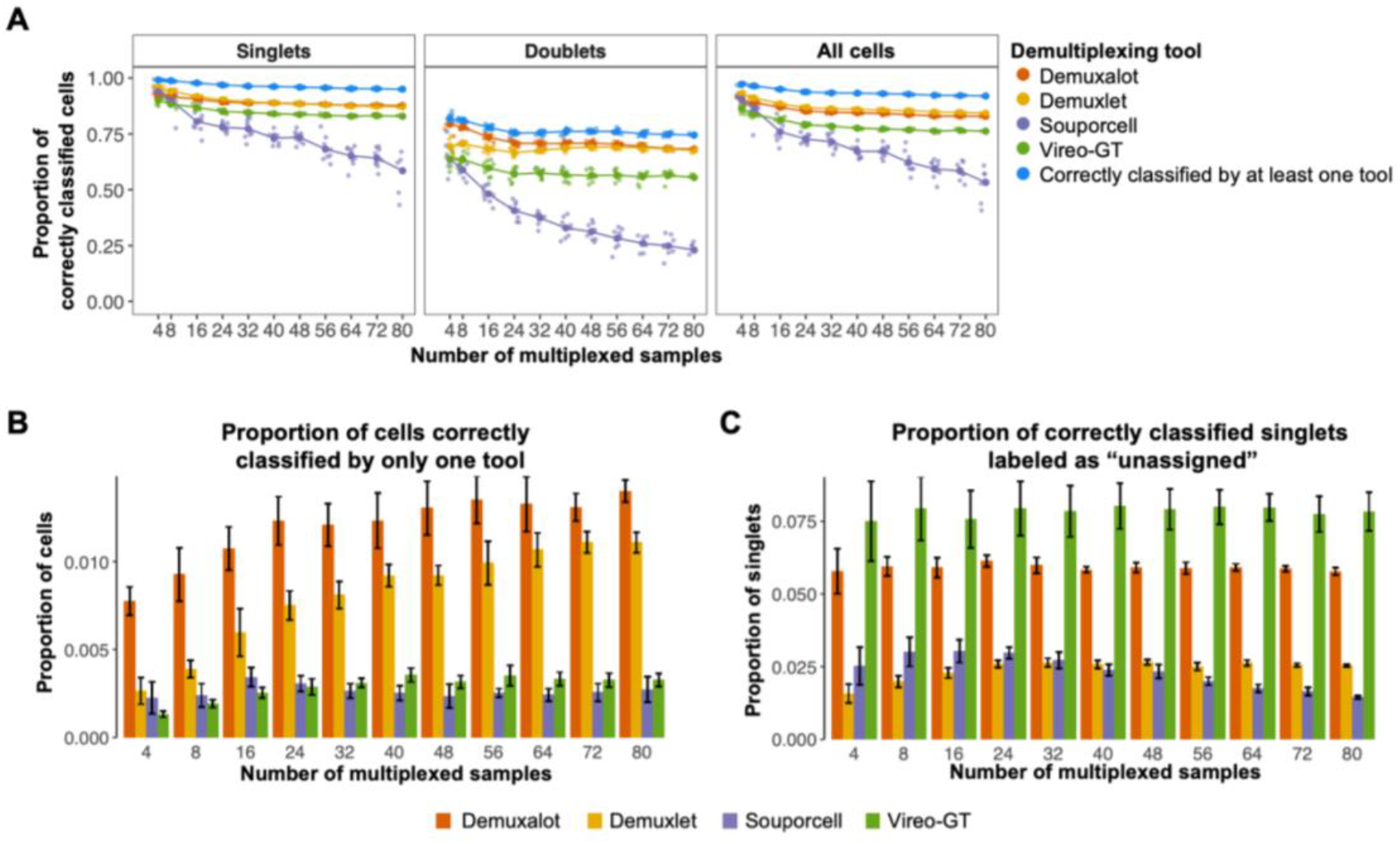
Evaluation of existing individual genetic demultiplexing tools. Evaluation of genetic demultiplexing tools with prior genotype information on 96 *in silico* pools with known ground-truth sample labels ranging in size from 4 to 80 multiplexed induced pluripotent stem cell (iPSC) lines from genetically distinct individuals, averaging 17,396 cells per pool and a 15% doublet rate. **A)** Line graphs showing the proportion of correctly classified singlets, doublets, and all cells by each individual genetic demultiplexing tool across varying numbers of multiplexed iPSC lines in a single pool (sample number). The large dots show the mean proportion of correct classifications by an individual tool across replicates at a given sample size (n = 9 per pool size). The blue points show the proportion of cells that were correctly classified by at least one individual genetic demultiplexing tool: Demuxalot, Demuxlet, Souporcell, or Vireo-GT. **B)** Bar chart showing the mean proportion of total cells from an individual pool correctly classified by only one genetic demultiplexing tool. Error bars represent one standard deviation from the mean. (n = 9 per pool size) **C)** Bar chart showing the proportion of correctly classified singlet cells labelled as “unassigned” (ambiguous singlet assignments) due to assignment probabilities below the recommended threshold of the respective genetic demultiplexing tool. Error bars represent one standard deviation from the mean. (n = 9 per pool size).

We also investigated the proportion of cells that were correctly classified by at least one genetic demultiplexing tool to designate the best possible performance of an ensemble method that successfully incorporates every correct classification from its constituent tools (**Figure 1A**). Across the 96 *in silico* pools, an average of 93.64% of cells were correctly classified by at least one tool. In comparison, Demuxlet, which demonstrated the best overall performance amongst individual tools, correctly classified 86.73% of cells, on average. Demuxalot was consistently responsible for the highest proportion of cells correctly classified by only one tool; 1.21% of pooled cells, on average, were correctly classified by Demuxalot only, followed by Demuxlet (mean = 0.83%), Vireo-GT (mean = 0.29%), and Souporcell (mean = 0.26%) (**Figures 1B; Additional File 1: Figure S1**). Conversely, a consensus framework, correctly classified only 81.06% of cells, on average (data not shown). Based on these results, we reasoned that an ensemble genetic demultiplexing method that can identify the most probable sample label from its constituent tools, independent of a consensus assignment, would increase the yield of correctly classified cells.

Next, we explored the frequency at which correctly classified singlets were labelled as unassigned because their assignment probability failed to meet the tool’s recommended probability threshold. Across the 96 *in silico* pools, Vireo-GT consistently showed the highest proportion of correctly classified singlets with insufficient assignment probabilities (Vireo-GT mean = 7.86%) followed by Demuxalot (mean = 5.91%), Demuxlet (mean = 2.44%) and Souporcell (mean = 2.34%) (**Figure 1C**). While a stringent probability threshold is important to prevent erroneous classifications in downstream analyses, we reasoned that the unnecessary removal of correctly classified cells could be mitigated by a carefully calibrated ensemble method that allocates additional assignment confidence to cells with matching sample labels across constituent tools, despite low internal tool-specific assignment probabilities.

We repeated the above analyses using the same 96 computationally multiplexed pools and the genetic demultiplexing tools that do not require prior genotype information: Freemuxlet, Souporcell, and Vireo. Here, we observed the same overarching limitations as when demultiplexing with prior genotype information: 1) decreasing demultiplexing performance as the number of multiplexed samples increased; 2) poor doublet detection performance compared to singlet classification; 3) high rates of cells only correctly classified by a single tool; and 4) discarded correctly classified cells due to insufficient assignment probabilities (**Additional File 1: Figure S2)**. When we compared demultiplexing with and without prior genotype information, we observed a trend towards a higher proportion of cells being correctly classified when prior genotype information was available, as previously seen in separate benchmarking analyses (9) (**Additional File 1: Figure S3**).

### Validating the Ensemblex framework on pools with known ground-truth sample labels

To mitigate the limitations of the individual genetic demultiplexing tools and maximize the analytic potential of multiplexed scRNAseq datasets, we developed Ensemblex (**Figure 2A)**. The Ensemblex workflow begins by demultiplexing pooled samples with four distinct demultiplexing algorithms, followed by three steps: 1) accuracy-weighted probabilistic ensemble; 2) graph-based doublet detection; and 3) ensemble-independent doublet detection **(Figure 2B)**. As output, Ensemblex returns its own cell-specific sample labels and corresponding assignment probabilities, as well as the sample labels and corresponding assignment probabilities for each of its constituent tools.

**Figure 2.**
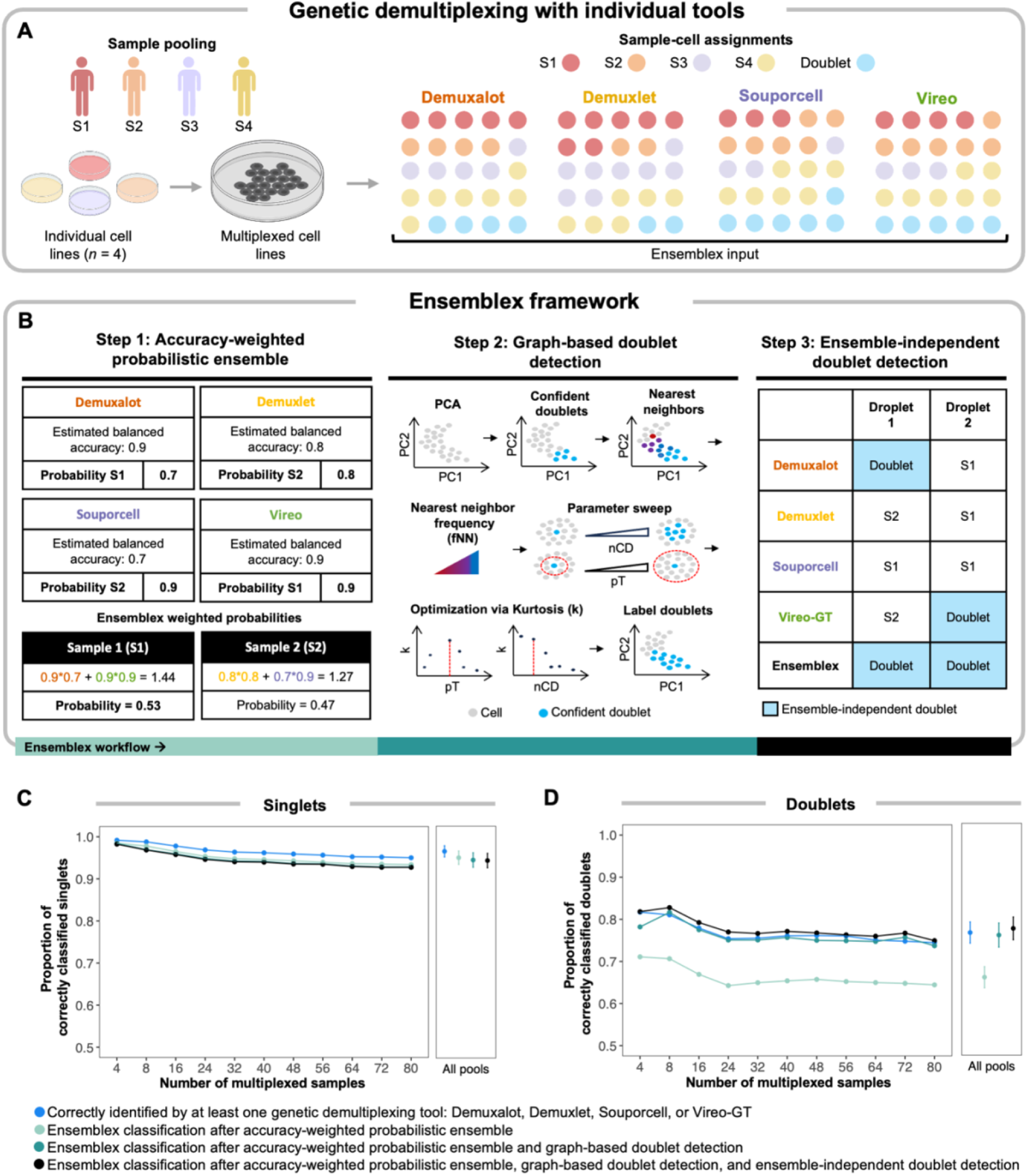
Characterization of the Ensemblex framework. Ensemblex is a probabilistic-weighted ensemble genetic demultiplexing framework for single-cell RNA sequencing analysis, which was designed to leverage the most probable sample labels from each of its constituent tools: Demuxalot, Demuxlet, Souporcell, and Vireo when using prior genotype information or Demuxalot, Freemuxlet, Souporcell, and Vireo when prior genotype information is not available. **A)** The Ensemblex workflow begins with demultiplexing pooled cells from genetically distinct individuals by each of the constituent tools. The outputs from each individual demultiplexing tool are then used as input into the Ensemblex framework. **B)** The Ensemblex framework comprises three distinct steps that are assembled into a pipeline: 1) accuracy-weighted probabilistic ensemble, 2) graph-based doublet detection, and 3) ensemble-independent doublet detection. **C-D)** Line graphs showng the contribution of each step of the Ensemblex framework on 96 *in silico* pools with known ground-truth sample labels ranging in size from 4 to 80 multiplexed induced pluripotent stem cell (iPSC) lines from genetically distinct individuals, averaging 17,396 cells per pool and a 15% doublet rate. The average proportion of correctly classified **C)** singlets and **D)** doublets across replicates at a given pool size is shown after sequentially applying each step of the Ensemblex framework with prior genotype information (n = 9 per pool size). The right panels show the average proportion of correct classifications across all 96 pools; error bars represent one standard deviation from the mean. The blue points show the proportion of cells that were correctly classified by at least one individual genetic demultiplexing tool: Demuxalot, Demuxlet, Souporcell, or Vireo-GT.

In response to our observation that certain cells are correctly classified by only one tool, we implemented the accuracy-weighted probabilistic ensemble component (Step 1) of the Ensemblex framework. In brief, this unsupervised weighting model identifies the most probable sample label for each cell by assigning weights to each tool’s assignment probabilities based on their estimated balanced accuracy for the dataset (see “Methods”) (**Figures 2B)** (16). Ensemblex then retains the sample label with the highest cumulative probability across its constituents. However, one challenge for this framework is computing the balanced accuracy of the constituent tools for experimentally multiplexed pools that lack ground-truth labels. Therefore, to estimate the balanced accuracy of a particular constituent tool (e.g., Demuxalot) without ground-truth labels, Ensemblex leverages the cells with a consensus assignment across the three remaining tools (e.g., Demuxlet, Souporcell, and Vireo-GT) as a proxy for ground-truth. To validate this approach, we utilized *in silico* pools with known ground truth sample labels to compute the Adjusted Rand Index (ARI) between Ensemblex’s sample labels when the balanced accuracy of the constituent tools was computed using consensus labels or ground-truth labels. Here, we consistently observed a mean ARI > 0.99, independent of the number of multiplexed samples in a pool, suggesting high assignment concordance between the two approaches (**Additional File 1: Figure S4)**. Applying the accuracy-weighted probabilistic ensemble component to the 96 *in silico* pools correctly classified 94.98% of singlets, on average, across all pools, approaching the number of singlets that were correctly classified by at least one constituent tool (mean = 96.48%) (**Figure 2C**). In contrast, only 66.01% of doublets, on average, were correctly identified across all pools after Step 1, compared to 76.59% of doublets that were correctly identified by at least one constituent tool (**Figure 2D**).

Given that previous analyses have demonstrated strong doublet call discordance across genetic demultiplexing tools (10), it was unsurprising that Step 1 of the Ensemblex framework performed poorly on doublet identification. Therefore, instead of relying on the cell type classifications of the constituent tools (i.e., singlet or doublet), we elected to leverage the doublet-related features (e.g., doublet probability; see “Methods”) returned by the constituent tools to identify the cells with the highest doublet likelihood, independent of the existing classifications. We implemented this approach in the graph-based doublet detection component (Step 2) of the Ensemblex framework, which was specifically designed to increase the rate of true doublet detection. Step 2 begins by identifying the top *n* most confident doublets in the pool (see “Methods”). Then, based on the Euclidean distances in principal component analysis (PCA) space, the cells that appear most frequently amongst the nearest-neighbors of the high confident doublets and exceed the optimized percentile threshold for the nearest-neighbor frequency are labelled as doublets by Ensemblex (**Figure 2B; Additional File 1: Figure S5;** see “Methods”). Upon applying the graph-based doublet detection component to the 96 *in silico* pools following Step 1, Ensemblex correctly identified 76.00% of doublets, on average: a 9.99% increase in doublet detection from Step 1. In turn, the average proportion of correctly classified singlets across all pools (94.43%) decreased by only 0.55% (**Figure 2D**).

The ensemble-independent doublet detection component (Step 3) of the Ensemblex framework was implemented to further improve doublet detection. Step 3 was motivated by our observation that certain tools, namely Demuxalot and Vireo, showed high doublet detection specificity (mean = 0.99) on *in silico* pools with known ground-truth sample labels, but that Steps 1 and 2 failed to incorporate a subset of these correct doublet calls (**Additional File 1: Figure S6**). Therefore, by default, Ensemblex accepts the doublet calls made by Demuxalot and Vireo-GT (**Figure 2B**). Applying the ensemble-independent doublet detection component to the 96 *in silico* pools following Steps 1 and 2 further increased the average proportion of correctly identified doublets across all pools by 1.58% for a total of 77.63% of doublets detected, while only decreasing the average proportion of correctly classified singlets by 0.13% for a total of 94.30% of singlets correctly classified (**Figures 2C and 2D**). Notably, owing to the graph-based doublet detection component, the average proportion of doublets identified by Ensemblex exceeded the average proportion of doublets that were correctly classified by at least one constituent tool.

While the three-step workflow of the Ensemblex pipeline was designed to maximize the balance between singlet classification and doublet identification, we do prioritize the identification of doublets at the expense of a slightly lower singlet yield to minimize technical noise in the data. However, we recognize that different experimental designs will require varying levels of doublet detection stringency; thus, users can modify the percentile thresholds for graph-based doublet detection and nominate different tools for ensemble-independent doublet detection (see “Methods”).

### Benchmarking Ensemblex on pools with known ground-truth sample labels

To benchmark Ensemblex against Demuxalot, Demuxlet, Souporcell, and Vireo-GT with prior genotype information, we first utilized the 96 *in silico* pools with known ground-truth sample labels to assess how Ensemblex’s demultiplexing performance varied as the number of multiplexed samples approached a cohort scale (4-80 samples). Unlike doublets, singlets were only considered correctly classified if their assignment probability exceeded the recommended threshold of the respective tool. On average across all pools, Ensemblex showed a higher proportion of correctly classified singlets (mean = 92.19%), doublets (mean = 77.63%), and all cells (mean = 90.12%) than the other tools. In comparison, Demuxlet, widely considered the “gold standard” tool, correctly classified 89.72% of singlets, 68.57% of doublets, and 86.73% of all cells, on average (**Figures 3A-3C**). Importantly, the discrepancy in the proportion of correctly classified cells between Ensemblex and the next-best tool was amplified as the number of multiplexed samples increased from 4 (2.78%) to 80 (3.52%), demonstrating that our ensemble method was able to partially mitigate decreased demultiplexing accuracy as the pools approach a population scale.

**Figure 3.**
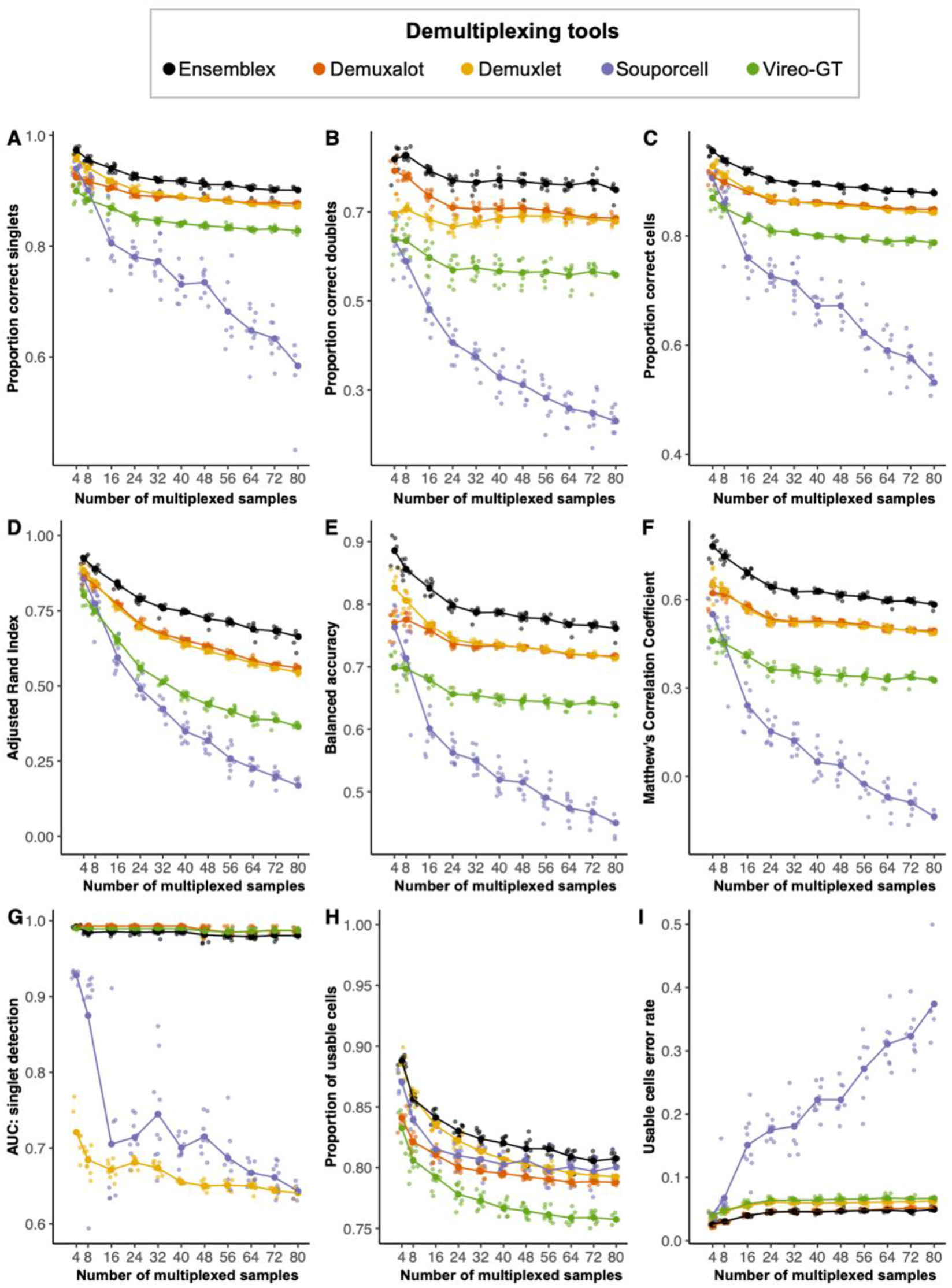
Ensemblex ground-truth benchmarking on computationally multiplexed pools. The genetic demultiplexing tools with prior genotype information were evaluated on 96 *in silico* pools with known ground-truth sample labels ranging in size from 4 to 80 multiplexed induced pluripotent stem cell (iPSC) lines from genetically distinct individuals, averaging 17,396 cells per pool and a 15% doublet rate. A singlet was considered correctly classified if the assigned sample label matched the ground-truth sample label and the assignment probability exceeded the recommended threshold for the respective tool; a doublet was considered correctly classified if the assigned sample label matched the ground-truth sample label, regardless of the assignment probability. **A-I)** Line graphs showing the performance of Ensemblex and the individual genetic demultiplexing tools across evaluation metrics. The large dots show the mean value for each tool across replicates at a given sample size (n = 9 per pool size). **A)** Proportion of correctly classified singlets. **B)** Proportion of correctly classified doublets. **C)** Proportion of correctly classified cells. **D)** Adjusted Rand Index between each tool’s sample labels and the ground-truth sample labels. **E)** Balanced accuracy of each tool. **F)** Matthew’s Correlation Coefficient of each tool. **G)** Area under the receiver operating characteristic curve (AUC) of the singlet assignment probability for each tool. **H)** Proportion of usable cells returned by each tool. Usable cells were defined as cells classified by singlets with an assignment probability exceeding the recommended threshold of the respective tool. **I)** Error rate amongst the usable cells returned by each tool; erroneous classifications comprised of true doublets labeled as singlets or true singlets assigned to the wrong sample.

Next, we applied evaluation metrics for classification models to gauge the overall performance of the genetic demultiplexing tools. We first computed the ARI to evaluate the similarity between the demultiplexed sample labels and the ground-truth sample labels. Here, Ensemblex showed the highest ARI with the ground truth sample labels across all pools (mean = 0.76), followed by Demuxalot (mean = 0.67) and Demuxlet (mean = 0.66) (**Figure 3D**). We then computed the balanced accuracy to evaluate the binary classification performance — singlet or doublet — of each genetic demultiplexing tool as well as the Matthew’s Correlation Coefficient (MCC), which previous work has suggested is more reliable and informative for classification cases where positive (singlet) and negative (doublet) cases have the same analytic importance (17). Across all pools, Ensemblex showed the highest balanced accuracy (mean = 0.80) and MCC (mean = 0.64), whereas Demuxalot and Demuxlet showed average balanced accuracies of 0.74 and 0.75, respectively, and both tools showed an average MCC of 0.54 (**Figures 3E and 3F**). To evaluate how well Ensemblex’s confidence score (see “Methods”) and each constituent tool’s assignment probability corresponded to the accuracy of their singlet classification, we plotted the area under the receiver operating characteristic curve (AUC). Although Demuxalot (mean = 0.99) and Vireo-GT (mean = 0.99) showed the highest AUC across all pools on average, Ensemblex’s AUC was comparable (mean = 0.98) (**Figure 3G**).

Finally, we investigated the proportion of usable cells returned by each demultiplexing tool and the error rate amongst usable cells. We define usable cells as singlet classifications exceeding the recommended probability threshold of the respective tool, while the error rate amongst usable cells constituted incorrectly classified singlets to the wrong donor-of-origin or true doublets incorrectly classified as singlets. We observed that, on average, Ensemblex returned the highest proportion of usable cells across all pools (82.66%), followed by Demuxlet (81.66%), Souporcell (81.01%), Demuxalot (79.99%), and Vireo-GT (77.53%) (**Figure 3H**). Importantly, Ensemblex showed the lowest error rate amongst usable cells (4.34%), followed by Demuxalot (4.43%), Demuxlet (5.77%), Vireo-GT (6.16%), and Souporcell (21.82%) (**Figure 3I**).

Using computationally multiplexed pools comprised of 24 iPSC lines, we further assessed how the performance of Ensemblex varied as a function of the number cells in a pool when prior genotype information was available. Here, we observed that our ensemble method consistently outperformed the individual demultiplexing tools (**Additional File 1: Figure S7**). When cells are pooled experimentally, it is reasonable to expect some iPSC lines to be underrepresented in the pool. Therefore, to assess Ensemblex’s demultiplexing performance in the presence of an underrepresented iPSC line, we produced computationally multiplexed pools comprising of 24 samples, with one sample showing varying degrees of under representation. Again, we observed that Ensemblex consistently outperformed the individual tools (**Additional File 1: Figure S8**).

Finally, we repeated the above analyses to assess whether the benefits of using Ensemblex to demultiplex with prior genotype information extended to cases where prior genotype information is not available. In doing so, we observed a trend towards better overall performance by Ensemblex; however, the discrepancy between Ensemblex and the top-performing individual tools, namely Freemuxlet and Souporcell, was less pronounced than when demultiplexing with prior genotype information (**Additional File 1: Figures S9-S11).**

Taken together, these results indicate that the Ensemblex framework mitigates the limitations of the individual tools, leading to greater overall demultiplexing performance across computationally multiplexed pools with known ground-truth labels. Ultimately, Ensemblex’s improved demultiplexing performance translates to a higher recovery of usable cells for downstream analyses as well as a higher accuracy amongst usable cells, limiting the unnecessary removal of cells from the dataset and mitigating the introduction of technical artifacts into biological analyses.

### Evaluating Ensemblex on experimentally pooled samples with donor-specific oligonucleotide labels

To determine whether Ensemblex’s improved performance across the *in silico* pools is reflected in real-world multiplexed experiments, we applied Ensemblex to an experimentally multiplexed pool composed of NSCLC dissociated tumor cells from 7 donors, hereafter referred to as the NSCLC dataset (18). Importantly, these NSCLC cells were labelled with donor-specific Cell Multiplexing Oligonucleotides (CMOs), providing a proxy for ground-truth sample labels to evaluate the performance of the genetic demultiplexing tools. For this experiment, we used HTOdemux (19) to assign the cells back to their donor-of-origin based on the CMO expression profiles. HTOdemux confidently assigned 19,695 cells, of which 15,534 (78.87%) were assigned to individual donors and 4,161 (21.13%) were assigned as doublets; 769 cells (3.76%) were unassignable at a positive quantile of 0.99 and were excluded from downstream analyses (**Figures 4A)**. Application of the Ensemblex framework with prior genotype information to the NSCLC dataset achieved a singlet true positive (TP) rate of 96.92% and doublet TP rate of 66.21% (**Figure 4B**). To evaluate the benefits of utilizing the entire Ensemblex workflow (Steps 1-3), we investigated the contribution of each step of the Ensemblex framework to the overall demultiplexing accuracy. Applying graph-based doublet detection (Step 2) and ensemble-independent doublet detection (Step 3) to the accuracy weighted assignments obtained from Step 1 increased the proportion of correctly identified doublets by 14%, while slightly decreasing the proportion of correctly classified singlets by 0.05% (**Additional File 1: Table S1**). Although users can elect to utilize different step-combinations of the Ensemblex pipeline, these results reaffirm that leveraging the entire workflow maximizes the overall demultiplexing accuracy by achieving a meticulous balance between singlet classification and doublet identification.

**Figure 4.**
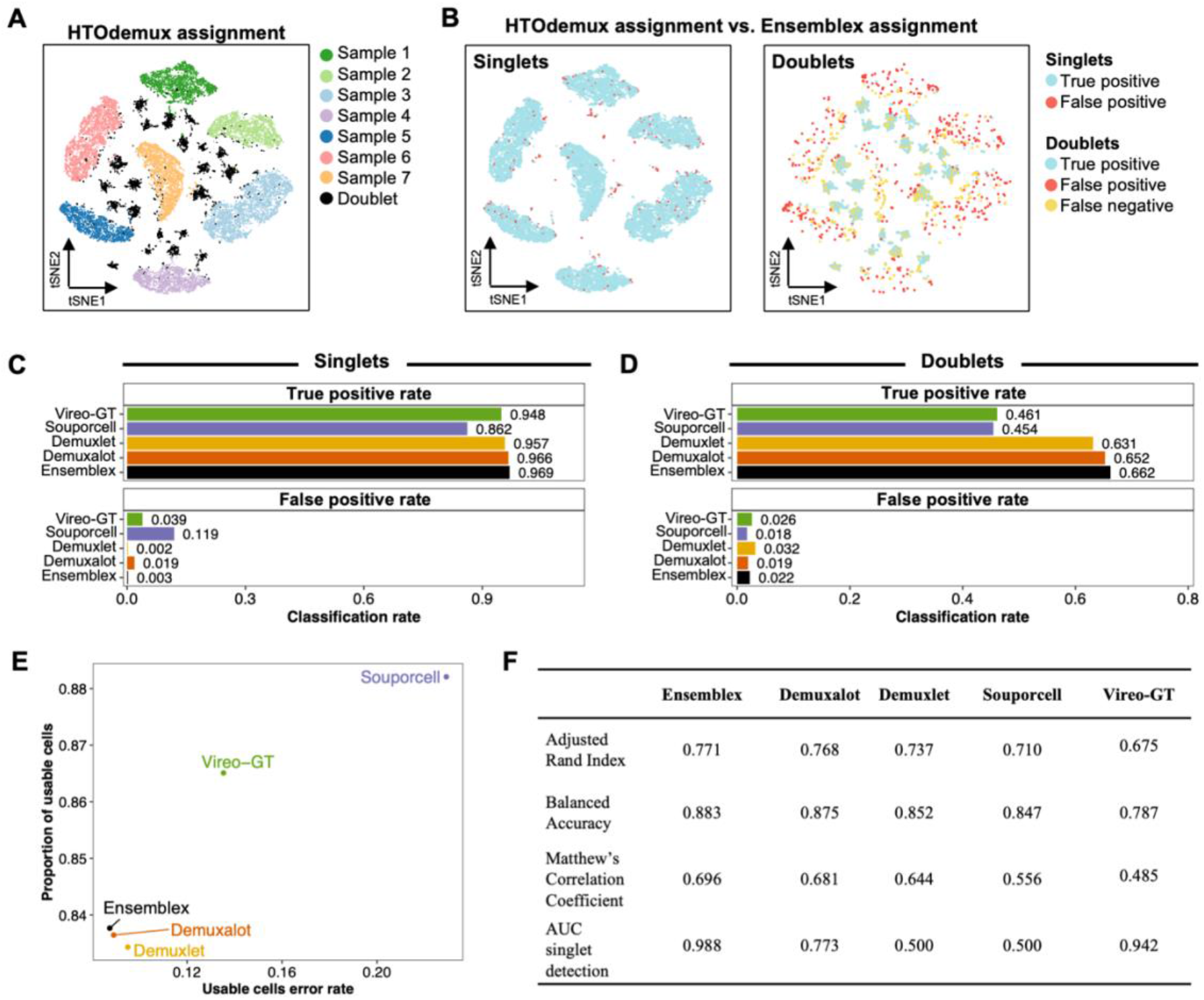
Evaluating Ensemblex on experimentally multiplexed cells using donor-specific oligonucleotide labels as a proxy for ground-truth. Non-small cell lung cancer (NSCLC) dissociated tumor cells from 7 individuals were pooled and labelled with donor-specific oligonucleotide-labels. Cells were demultiplexed according to their expression of donor-specific oligonucleotide labels by HTOdemux; HTOdemux’s sample labels were used as a proxy for ground truth. True positives (TP) singlets were defined as cells classified as singlets by both HTOdemux and Ensemblex with matching sample labels; false positives (FP) singlets were defined as cells classified as singlets by both HTOdemux and Ensemblex but assigned to different donors. TP doublets were defined as cells classified as doublets by both HTOdemux and Ensemblex; FP doublets were defined as cells classified as singlets by HTOdemux and doublets by Ensemblex; false negatives (FN) doublets were defined as cells classified as doublets by HTOdemux and singlets by Ensemblex. **A)** T-distributed Stochastic Neighbor Embedding (t-SNE) visualization of HTOdemux’s sample labels. **B)** T-SNE visualization of Ensemblex’s demultiplexing performance using HTOdemux’s sample labels as ground truth for singlets (left) and doublets (right). **C)** Bar plots showing the singlet TP and FP rates for each genetic demultiplexing tool using HTOdemux’s sample labels as ground truth. **D)** Bar plots showing the doublet TP and FP rates for each genetic demultiplexing tool using HTOdemux’s sample labels as ground truth. **E)** Scatter plot showing the proportion of usable cells (confidently classified singlets) and the corresponding usable cell error rate for each genetic demultiplexing tool. **F)** Adjusted Rand Index, balanced accuracy, Matthew’s Correlation Coefficient, and area under the receiver operating characteristic curve (AUC) of the singlet assignment probability for each genetic demultiplexing tool.

Upon comparing Ensemblex’s demultiplexing performance with prior genotype information on the NSCLC dataset to the individual genetic demultiplexing tools, it emerged that our ensemble method obtained the highest singlet and doublet TP rates (**Figures 4C and 4D**). Ensemblex and Demuxlet also showed the lowest singlet false positive (FP) rates (0.25% and 0.21%, respectively), indicating that singlets were least frequently assigned to the wrong donor-of-origin by these two methods compared to Demuxalot (1.87%), Vireo-GT (3.91%), and Souporcell (11.94%). Souporcell and Vireo-GT returned the highest proportion of usable cells (confidently classified singlets; 88.21% and 86.51%, respectively); albeit, at the expense of high usable cell error rates (22.91% and 13.53%, respectively) (**Figure 4E**). In turn, Ensemblex, Demuxalot, and Demuxlet showed lower error rates across the usable cells (8.75%, 8.91%, and 9.51%, respectively), amongst which Ensemblex returned the highest proportion of usable cells (83.77%) compared to Demuxalot (83.64%) and Demuxlet (83.43%). Here, the relatively high error rate amongst usable cells returned by each demultiplexing tool is attributed to true doublets classified as singlets. Finally, we computed the ARI, balanced accuracy, MCC, and AUC for singlet detection for each tool and observed that Ensemblex again outperformed the remaining tools (**Figure 4F**). We repeated the above analyses without prior genotype information and observed a similar trend towards better overall performance by Ensemblex (**Additional File 1: Table S2 and Figure S12**). Together, these results corroborate that Ensemblex’s improved performance on the *in silico* pools extends to experimentally multiplexed samples.

### Application of Ensemblex to experimentally pooled, highly multiplexed subjects

To evaluate Ensemblex’s demultiplexing performance on experimentally pooled, highly multiplexed scRNAseq datasets with prior genotype information, we used pools containing iPSC lines from 22 donors that were differentiated towards DaN by Jerber et al., hereafter referred to as the DaN dataset (12) (**Figure 5A**). To capture the transcriptional changes throughout neurogenesis, Jerber et al. performed scRNAseq of the iPSC lines grown in pooled cultures at days 11, 30, and 52 of differentiation (**Figure 5A**). Using three technical replicates from each timepoint, we obtained 84,746 cells after performing quality control as previously described (12) (**Additional File 1: Table S3**). Each technical replicate was demultiplexed independently by Ensemblex and its constituent tools.

**Figure 5.**
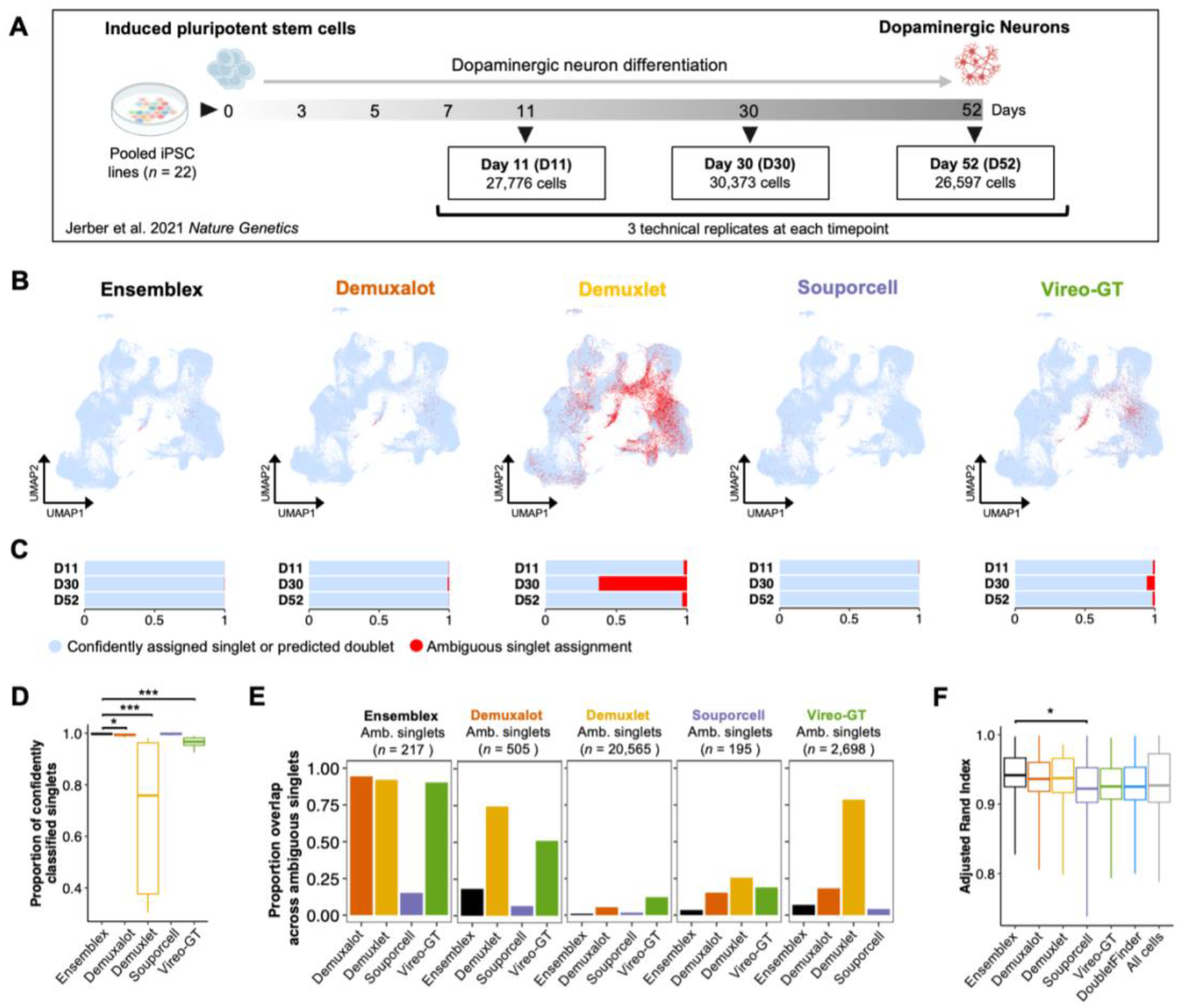
Application of Ensemblex to highly multiplexed, experimentally pooled cultures of differentiated dopaminergic neurons. A) Time line of iPSC pooling, dopaminergic neuron (DaN) differentiation, and sample collection from the DaN dataset by Jerber et al. (12). Three technical replicates at each time point (days 11, 30 and, 52 of differentiation) from pools containing 22 individual iPSC lines were used in the analysis. Across all timepoints and technical replicates, 84,746 cells were obtained for analysis. **B)** Uniform manifold approximation and projection (UMAP) plots showing confidently assigned singlets or predicted doublets (blue) and ambiguous singlets (singlet assignments with insufficient assignment probabilities; red) returned by each demultiplexing tool. **C)** Stacked bar chart showing the proportion of confidently assigned singlets or predicted doublets (blue) and ambiguous singlets (red) across technical replicates at each time point returned by each demultiplexing tool. **D)** Boxplot showing the proportion of confidently classified singlets across technical replicates and time points by each demultiplexing tool. Wilcoxon rank-sum tests were used to compare the proportion of confidently classified singlets by Ensemblex to that of its constituents (n = 9 pools). **E)** Bar chart showing the proportion of overlapping ambiguous singlet assignments amongst demultiplexing tools across technical replicates and time points (n = 9 pools). **F)** Boxplot showing the Adjusted Rand Index (ARI) assessing cluster stability across a range of 11 clustering resolutions (*n* clustering iterations = 25) after removing doublets identified by each demultiplexing tool. Wilcoxon rank-sum tests were used to compare the clustering ARI after removing Ensemblex doublets to the clustering ARI after removing doublets identified by each constituent tool. * Adjusted P-value < 0.05; ** adjusted P-value < 0.01; *** adjusted P-value < 0.001

To characterize the relationship between Ensemblex and its constituent demultiplexing tools, we computed the ARI between Ensemblex’s sample labels and those of its constituent as well as the percent contribution of each tool to Ensemblex’s final sample labels (**Table 2**). Notably, we observed that across day 30 technical replicates Demuxlet showed an ARI of 0.063 with Ensemblex and only contributed 29.74% to Ensemblex’s final sample labels. In contrast, across day 11 and 52 technical replicates Demuxlet showed an ARI of 0.928 and 0.884, respectively, and contributed 95.91% and 90.55%, respectively, to Ensemblex’s final sample labels. Importantly, Demuxlet’s variable contribution to Ensemblex’s sample labels across sequencing time points demonstrates our ensemble method’s ability to adapt to the relative performance of its constituent tools and override the classifications of a poorly performing tool on the respective dataset.

**Table 2.**
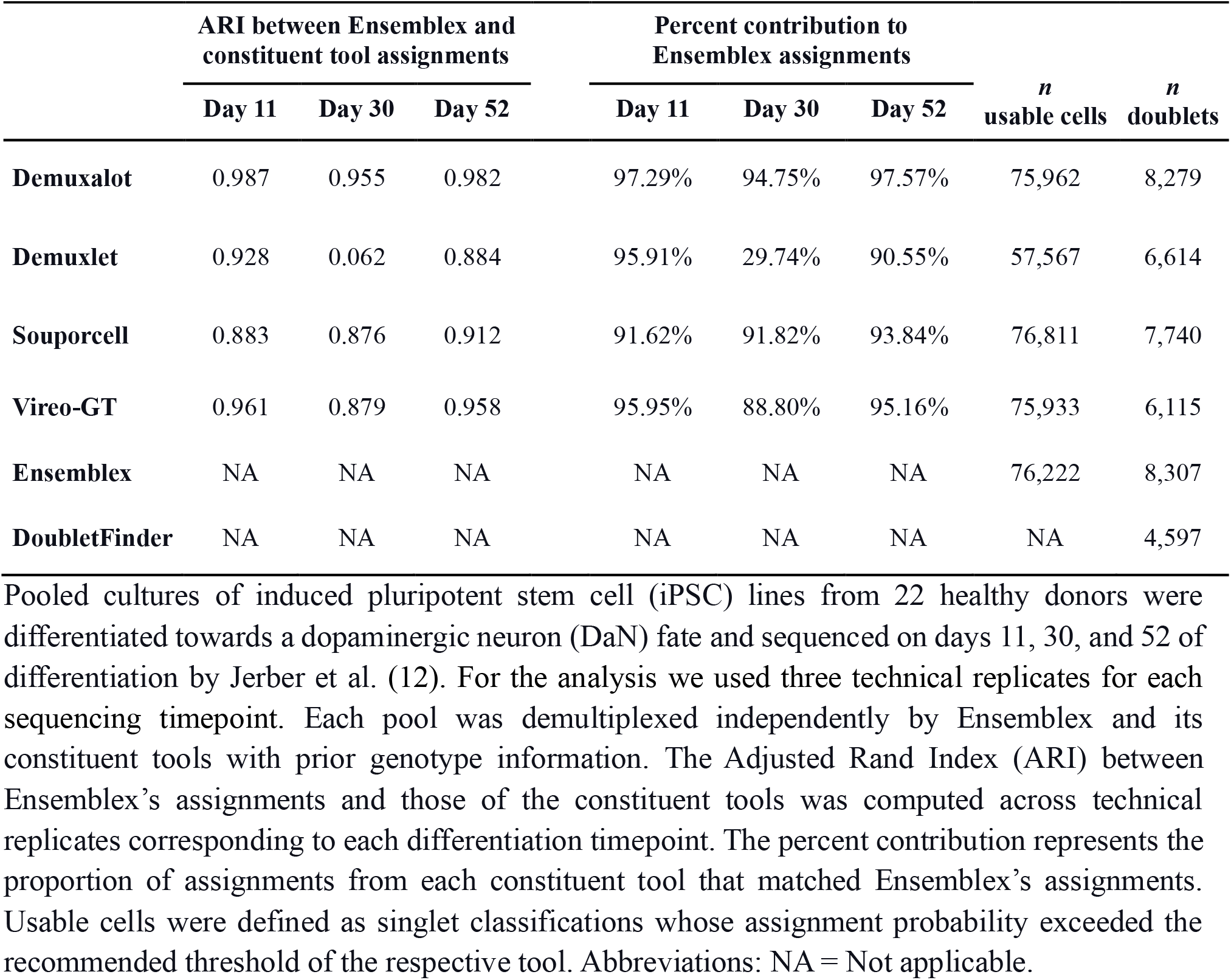
Application of Ensemblex to pooled cultures of dopaminergic neurons from 22 healthy controls.

To elucidate the discrepancy in Demuxlet’s contribution to Ensemblex’s sample labels across sequencing time points, we investigated the proportion of ambiguous singlet assignments from Ensemblex and its constituents. Ambiguous singlets are defined as singlet classifications whose assignment probabilities failed to meet the recommended threshold of the respective tool, leaving the identity of the pooled cell unresolved. Across 84,746 cells, Souporcell (195 singlets; 0.23% of cells) and Ensemblex (217 singlets; 0.26% of cells) showed the lowest proportion of ambiguous singlet assignments, followed by Demuxalot (505 singlets; 0.60% of cells) and Vireo-GT (2,698 singlets; 3.18% of cells). Strikingly, Demuxlet showed 20,565 ambiguous singlet assignments (24.27% of cells), with 92.04% derived from day 30 technical replicates, reflecting Demuxlet’s remarkably low contribution to Ensemblex’s sample labels for cells sequenced at this timepoint (**Figures 5B and 5C**). In accordance with previous analyses (9, 10), Demuxlet was consistently amongst the top performing constituent tools throughout our benchmarking analyses. Yet, its poor performance across day 30 technical replicates illustrates how the accuracy of individual tools can vary greatly between datasets, highlighting the importance of utilizing multiple distinct algorithms for genetic demultiplexing. We compared the mean proportion of confidently classified singlets across technical replicates from each time point (*n = 9)* between Ensemblex (99.72%) and each constituent demultiplexing tool using a Wilcoxon rank-sum test. After correction for multiple hypothesis testing, we observed that the mean proportion of confidently classified singlets by Ensemblex was significantly higher than Demuxalot (mean = 99.36%, P-value = 3.55e-3), Demuxlet (mean = 75.82%, P-value = 1.55e-5), and Vireo-GT (mean = 96.71%, P-value = 1.55e-5) (**Figure 5D**). Thus, despite Demuxlet’s unusually poor performance across day 30 technical replicates, Ensemblex still confidently classified 27,520 singlets (99.61% of singlet assignments) from these pools. Indeed, our ensemble method mitigates the consequences of a poorly performing constituent tool by outweighing the erroneous classifications. In contrast, using a consensus framework returned only 7,446 confidently classified singlets from day 30 technical replicates (20,074 fewer cells than Ensemblex), limiting the availability of data for downstream analyses.

To further evaluate the ambiguity amongst singlet classification, we investigated the intersection of ambiguous singlets across demultiplexing tools, reasoning that cells that are most challenging to demultiplex would be labelled as ambiguous across all tools (**Figure 5E**). The singlets that were assigned as ambiguous by Ensemblex showed the highest ambiguous singlet rate across the remaining tools (mean across all constituent tools = 73.04%; mean across Demuxalot, Demuxlet, and Vireo-GT = 92.32%). In contrast, while Souporcell showed the lowest ambiguous singlet rate overall, only 15.90% of its unassigned singlets, on average, were ambiguous across the remaining tools. These results indicate that the cells labelled as ambiguous by Ensemblex represent the cells that are most challenging to classify across the distinct demultiplexing algorithms. Indeed, limiting Ensemblex’s ambiguous singlet assignments to those that are most difficult to classify is critical for maintaining a balance between maximizing the number of usable cells and minimizing the introduction of technical artifacts into downstream analyses from misclassified cells.

Next, we compared the doublet predictions made by each genetic demultiplexing tool and DoubletFinder, a doublet detection tool that predicts doublets by estimating the similarity of the transcriptional profile of a pooled cell to artificial doublets generated by combining the transcriptional profiles of randomly selected cell pairs (20). Although the average number of unique molecular identifiers (UMI) per cell across doublets identified by each tool was significantly higher than the consensus singlets (**Additional File 1: Figure S13**), we observed a notable discrepancy in the number of doublets identified by each tool; DoubletFinder identified the fewest doublets (*n =* 4,597), while Ensemblex identified the most doublets (*n* = 8,307) (**Table 1**). Accordingly, all tools identified doublets that every other tool assigned as singlets (**Additional File 1: Figure S13**). While Ensemblex identified the highest number of doublets, it still returned a higher number of confidently classified singlets (*n =* 76,222) than Demuxalot (*n =* 75,962), Demuxlet (*n =* 57,567), and Vireo-GT (*n =* 75,933). Thus, even though the Ensemblex framework prioritizes the identification of doublets at the expense of a slightly lower singlet classification rate, our ensemble method still returns a high proportion of usable cells for downstream analyses.

To evaluate the impact of doublet removal on the stability of clusters in the DaN dataset, we performed 25 different random start iterations of the Louvain network detection at various clustering resolutions after removing the doublets identified by each tool (21). Removing the doublets identified by Ensemblex resulted in the highest ARI (mean ARI = 0.942), on average, across clustering resolutions (**Figure 5F**), suggesting the greatest cluster stability. However, Wilcoxon rank-sum tests only revealed a statistically significant difference in the cluster assignment ARI between Ensemblex and Souporcell (mean ARI = 0.922, P-value = 1.08e-2) after correction for multiple hypothesis testing. Nonetheless, the highest cluster stability after removal of Ensemblex’s putative doublets illustrates how improved doublet detection can translate to improved biological analyses and is reflective of its superior doublet identification performance on the benchmarking analyses.

### Evaluating the impact of demultiplexing tools on differential gene expression analysis

To evaluate the impact of genetic demultiplexing tools on scRNAseq DGE analysis, we multiplexed iPSC-derived NSCs from individuals with ADHD and controls (**Figure 6A**). NSCs were pooled and cultured until 100% confluence was reached. Two multiplexing experiments were performed: Experiment 1 (*n* ADHD = 7; *n* control = 6) and Experiment 2 (*n* ADHD = 9; *n* control = 7). After filtering cells for > 500 total and unique RNA transcripts, we obtained 30,433 cells across both pools. Louvain clustering on the integrated scRNAseq dataset identified 12 clusters, which were annotated as eight putative cell types (**Figure 6B**).

**Figure 6.**
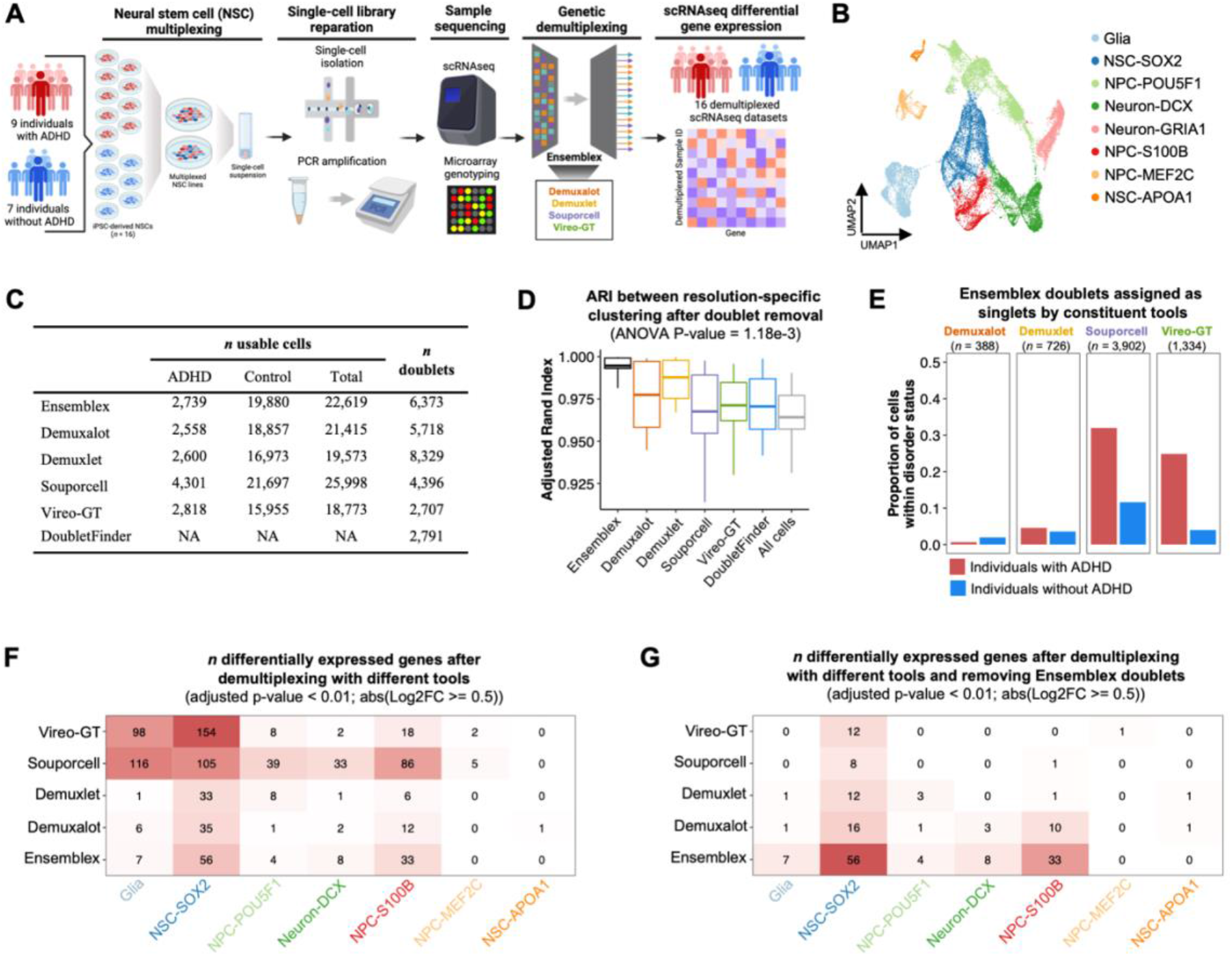
Evaluating the impact of discordant assignments between genetic demultiplexing tools on differential gene expression analysis. A) Schematic illustrating the workflow for the neural stem cell (NSC) dataset. Pooled induced pluripotent stem cell (iPSC)-derived neural stem cell cultures from individuals with attention deficit hyperactivity disorder (ADHD) and controls were collected in two separate experiments. NSCs were dissociated for single-cell RNA sequencing and prior genotype information of the pooled subjects was obtained through microarray genotyping. The pools were demultiplexed by Ensemblex and its constituents with prior genotype information and differential gene expression (DEG) was computed between ADHD and controls. **B)** Uniform manifold approximation and projection (UMAP) plot showing the putative cell types. **C)** Summary of the number of usable cells — singlets above the recommended probability threshold of the respective demultiplexing tool — assigned to ADHD donors and controls and the number of identified doublets by each demultiplexing tool. **D)** Boxplot showing the Adjusted Rand Index (ARI) assessing cluster stability across a range of 11 clustering resolutions (*n* clustering iterations = 25) after removing doublets identified by each demultiplexing tool. A one-way Analysis of Variance (ANOVA) test comparing the ARI after removing doublets identified by each tool revealed a significant difference between tools (n = 11 clustering resolutions; P-value = 1.18e-3). **E)** Proportion of ADHD and control cells identified as putative doublets by Ensemblex that were assigned as singlets by the constituent demultiplexing tools. **F)** Heatmap showing the number of cell-type specific DEGs between ADHD and controls using the subject labels of each demultiplexing tool. **G)** Heatmap showing the number of cell-type specific DEGs between ADHD and controls using the subject labels of each demultiplexing tool and removing putative doublets identified by Ensemblex. Cell-types not shown in the heatmaps had no DEGs passing the adjusted P-value < 0.01 and |Log2FC >= 0.5| threshold across all tools.

We independently demultiplexed both pools using Ensemblex and its constituents to assign the cells back to their donor-of-origin with prior genotype information (**Figure 6C).** The number of cells assigned to ADHD and control donors by each genetic demultiplexing tool is shown in **Additional File 1: Table S6**. Importantly, the NSC dataset provides a valuable illustration of the consequences of unnecessarily discarding cells from downstream analyses. For example, Ensemblex and Vireo-GT returned 2,387 and 882 confidently assigned *GRIA1*^high^ neurons, respectively, whereas a consensus approach would have confidently assigned only 563 *GRIA1*^high^ neurons (**Additional File 1: Table S6**).

Each genetic demultiplexing tool predicted the ADHD cells to be vastly underrepresented compared to the control cells; Ensemblex assigned 2,739 cells to individuals with ADHD and 19,880 cells to controls, suggesting that the ADHD iPSC lines were lost throughout the culturing and sequencing process (**Figure 6C).** Additionally, we observed a notable difference in the number of identified doublets across the tools; Vireo-GT identified the fewest doublets (*n* = 2,707), while Demuxlet identified the most doublets (*n* = 8,329) (**Figure 6C)**. We aimed to characterize the change in cluster stability after removing the doublets identified by each tool and observed that removing the doublets identified by Ensemblex resulted in the highest ARI (mean ARI = 0.995), on average, across clustering resolutions (**Figure 6D**). A one-way ANOVA test comparing the clustering ARI after removal of doublets identified by each tool revealed a significant difference between tools (P-value = 1.18e-3). Demuxlet (*n* = 8,329) identified more doublets than Ensemblex (n = 6,373), but exhibited lower cluster stability (ARI), suggesting that increased cluster stability is not merely representative of the number of doublets removed but rather the quality of doublet removal.

Given the underrepresentation of ADHD cells across the dataset, we elected to investigate the cells that were identified as doublets by Ensemblex but assigned as singlets by the constituent tools and how these putative doublets were distributed across samples according to disorder status. Demuxalot (*n =* 388) and Demuxlet (*n =* 726) assigned a relatively low number of Ensemblex’s doublets as singlets, which represented 0.66% and 4.58% of ADHD sample assignments, respectively, and 1.97% and 3.58% of control sample assignments, respectively (**Figure 6E**). In contrast, Souporcell (*n =* 3,902) and Vireo-GT (*n =* 1,334) assigned a relatively high number of Ensemblex’s doublets as singlets, which represented 31.97% and 24.88% of ADHD sample assignments, respectively, and 11.65% and 3.97% of control sample assignments, respectively, illustrating how variable doublet detection can impact the assembly of cells assigned to donor categories and which cells are retained for downstream analyses.

Finally, we used the model-based analysis of single-cell transcriptomics (MAST) statistical framework to compute cell-type specific DGE between individuals with ADHD and controls using the demultiplexed sample labels from each tool (22). We observed a significant discrepancy in the number of cell type-specific differentially expressed genes (DEGs; adjusted P-value < 0.01; absolute log2 fold change > 0.5) depending on the demultiplexing tool used (**Figure 6F**). Most notably, for glia cells Souporcell identified 116 DEGs; Vireo-GT identified 98 DEGs; Ensemblex identified 7 DEGs; Demuxalot identified 6 DEGs; and Demuxlet identified 1 DEG. Similar patterns were observed across *SOX2*^high^ NSCs, *POU5F1*^high^ neural progenitor cells (NPC), *S100B*^high^ NPCs, and *DCX*^high^ neurons, whereby Souporcell or Vireo-GT’s sample labels resulted in a remarkably high number of DEGs compared to Ensemblex, Demuxalot, and Demuxlet. Given that Souporcell and Vireo-GT made relatively few doublet calls and that 31.97% and 24.88% of ADHD sample assignments made by Souporcell and Vireo-GT, respectively, were putative doublets identified by Ensemblex, we elected to repeat the DGE analysis using the demultiplexed sample labels from each tool but this time we removed all putative doublets identified by Ensemblex. In doing so, we observed a decrease in the number of DEGs identified by Souporcell and Vireo-GT across cell types, suggesting that the putative doublets identified by Ensemblex, which were classified as singlets by Souporcell and Vireo-GT, were driving the initial signals (**Figure 6G**). For example, the number of glia-specific DEGs decreased from 116 to 0 with Souporcell’s sample labels, and 98 to 0 with Vireo-GT’s sample labels. Given that the NSC dataset lacked ground-truth sample labels, we could not definitively determine which cells were true doublets; however, the increase in clustering ARI after removal of Ensemblex’s putative doublets (**Figure 6D**), coupled with Ensemblex’s improved doublet identification performance on pools with known ground-truth sample labels (**Figure 2B**), afforded confidence to assume that our ensemble method performed favorably. Nonetheless, this analysis reveals that the choice of demultiplexing tool can greatly impact biological analyses.

## Conclusion

Multiplexing protocols, coupled with the introduction of genetic demultiplexing tools constituted a significant advancement for scRNAseq by providing a feasible means to dramatically increase the throughput of biological replicates. As the demand for population-scale scRNAseq analysis continues to grow with the maturation of singe-cell technologies, the prospect of multiplexing entire cohorts has emerged. However, the realization of this goal is impeded by the limitations of the current genetic demultiplexing tools. These include decreasing demultiplexing performance as the number of multiplexed samples increases (9, 10), relatively poor doublet detection performance (10), relatively high rates of cells that can only be correctly classified by single algorithms, the unnecessary removal of correctly classified cells due to insufficient assignment probabilities, and highly variable demultiplexing performance between datasets (10). In this work we presented Ensemblex, which offers a unique solution to these limitations by meticulously implementing distinct demultiplexing algorithms into a robust, accuracy-weighted ensemble framework that is exceptionally equipped to classify highly multiplexed pools.

We applied Ensemblex to a diverse array of computationally and experimentally multiplexed scRNAseq datasets. Benchmarking analyses on pools with known ground-truth sample labels revealed Ensemblex’s superior demultiplexing performance across pools reaching 80 multiplexed samples, which translated to a higher proportion of cells retained for downstream analyses and lower error rates amongst classified cells. Ensemblex also demonstrated a notable advancement for identifying heterogenic doublets, which is a well-documented limitation of the genetic demultiplexing tools currently available (9, 10, 15). While previous analyses indicated that the number of multiplexed samples in a pool directly impacted doublet detection efficiency (15), we showed that Ensemblex’s ability to identify doublets remained relatively constant when >24 samples were multiplexed. Our findings suggest that super loading cells prior to sequencing — which will result in a higher number of usable cells but a higher a doublet rate (6) — followed by heterogenic doublet detection by Ensemblex, may be a viable approach for implementing population-scale multiplexing in practice. We also demonstrated that the performance of individual genetic demultiplexing tools can be highly dataset-dependent, reflecting the findings of previous work (10). However, due to its unsupervised weighting model, we showed that Ensemblex is resistant to poorly performing constituent tools, maximizing the consistency of its demultiplexing performance. Nonetheless, if each constituent tool performs poorly on a given dataset, the poor performance will be reflected in Ensemblex’s demultiplexing accuracy. Finally, we illustrated that discordant sample assignments amongst genetic demultiplexing tools can greatly impact DGE analyses, necessitating that investigators carefully consider their choice of genetic demultiplexing tool. Although untested, we anticipate that the impacts of discordant sample assignments amongst genetic demultiplexing tools on biological interpretations would be exacerbated for computational analyses that consider the specific donor identity of the pooled cells, such as expression quantitative trait loci (eQTL) analyses, as opposed to donor groups (i.e., case and control). Due to Ensemblex’s ability to seamlessly integrate multiple algorithms into an adaptable framework, we argue that our ensemble method achieves unmatched reliability for experimentally multiplexed pools that lack ground truth sample labels.

Undoubtedly, a limitation of utilizing an ensemble method for genetic demultiplexing is the necessity to run each individual demultiplexing algorithm, which can be computationally expensive. Yet, in the absence of comparing demultiplexed sample labels across tools, poor performance by a given individual algorithm on experimentally multiplexed pools is undetectable, and the risk of introducing technical artifacts and losing usable cells for downstream analyses is prominent. As such, we believe that the relatively high computational cost of Ensemblex is a worthwhile investment to maximize the biological insight obtained from multiplexed scRNAseq datasets. To mitigate the burden of genetic demultiplexing by multiple individual tools, we provide a coherent pipeline that runs each constituent demultiplexing tool in parallel and seamlessly processes the respective output files with the Ensemblex algorithm.

Compared to when demultiplexing was informed by prior genetic data of the pooled samples, the improvement of Ensemblex over its constituent tools was far less pronounced for genotype-free demultiplexing cases. All demultiplexing tools, including Ensemblex, showed drops in demultiplexing performance when >16 samples were multiplexed in a pool without prior genotype information. Nonetheless, Ensemblex still constitutes an advancement over the individual tools for genotype-free demultiplexing cases due to the robustness achieved by incorporating distinct demultiplexing algorithms, which protects against the prospect of poorly performing individual tools on the respective dataset. Furthermore, an intrinsic limitation of demultiplexing without prior genotype information is that samples cannot be directly linked to metadata, leaving the sample identity of the inferred clusters unresolved (9). Although challenging, this limitation can be mitigated by identifying a small subset of discriminatory variants from the reconstructed genotypes of the constituent demultiplexing tools, which could be used to manually assign the computed clusters to samples if such discriminatory variants are known by the investigator. While the Ensemblex pipeline provides users the option to demultiplex pools with or without prior genotype information, we assert that users take caution when electing to perform population-scale multiplexing experiments without using prior genetic data.

Genetic demultiplexing tools have been used extensively for scRNAseq analysis across many disciplines in the biological sciences, including microbiology (8), model organisms (15), cancer biology (23), and neurodegenerative disease (12). Recent work has also evaluated the utility of genetic demultiplexing tools for different single-cell, read-based modalities such as single-nuclei RNA sequencing (snRNAseq) and single-nuclei assay for transposase-accessible chromatin sequencing (scATACseq) (24). Although untested, we expect Ensemblex to prove beneficial in demultiplexing for these assays, but comprehensive benchmarking with the appropriate datasets is required and was not explored here.

We expect numerous biological fields to exploit the benefits of Ensemblex through its application to highly multiplexed pools comprising cells from many genetically distinct individuals. Specifically for biomedical sciences, the preparation and labour costs of scRNAseq remains prohibitively expensive for analyzing entire cohorts of patients, which is critical for characterizing the genetic heterogeneity and etiological diversity of disease, and for maintaining sufficient statistical power for detecting associations between transcriptional changes and clinical or pathological observations (1). By increasing the throughput of biological replicates, multiplexing has rendered the prospect of analyzing entire patient cohorts with single-cell transcriptomics feasible. Highly-multiplexed scRNAseq experiments have already been presented in the literature and, to the best of our knowledge, have pooled up to 24 samples in a single dish (12). However, we demonstrated that Ensemblex’s demultiplexing accuracy remains relatively constant when >24 samples are multiplexed at concentrations that abide by the current limitations of experimental protocols, suggesting that Ensemblex equips the research community with the necessary computational framework to expand the upper limits of the number of genetically distinct individuals in a single pool.

While multiplexing mitigates the labour and consumable costs of scRNAseq analysis, the cost of sequencing remains expensive and the increasing number of genetically distinct individuals in a single pool necessitates that a greater number of cells must be sequenced to ensure adequate representation. Accordingly, Ensemblex is equipped to demultiplex pools comprising cells from more genetically distinct individuals than is feasible with the current laboratory technologies. However, we expect that the cost of sequencing will continue to decrease with the maturation of the technology, and our tool will be in place for when the anticipated wet lab advancements are realized. Overall, we conclude that Ensemblex constitutes a notable advancement towards the pressing demand for population-scale single-cell transcriptomics.

## Methods

### Ensemblex framework overview

Ensemblex is an ensemble genetic demultiplexing framework for scRNAseq sample pooling that was designed to identify the most probable sample labels from each of its constituent tools: Demuxalot (5), Demuxlet (6), Souporcell (8), and Vireo (9) when demultiplexing with prior genotype information or Demuxalot, Freemuxlet (6), Souporcell, and Vireo when demultiplexing without prior genotype information. After running each constituent demultiplexing tool in parallel, Ensemblex merges the output files containing the sample-cell assignments from each tool and performs three distinct steps of the Ensemblex pipeline:

1. Accuracy-weighted probabilistic ensemble;
2. Graph-based doublet detection;
3. Ensemble-independent doublet detection.

Upon obtaining the final Ensemblex sample labels (donor-of-origin identity of the pooled cells), the singlet assignment confidence score is computed.

### Step 1: Accuracy-weighted probabilistic ensemble

Ensemblex utilizes an unsupervised weighting model to identify the most probable sample label for each cell. Ensemblex weighs each constituent tool’s assignment probability distribution by its estimated balanced accuracy for the dataset in a framework adapted from the work of Large et al. (16). To estimate the balanced accuracy of a particular constituent tool (e.g., Demuxalot) for experimentally multiplexed datasets lacking ground-truth labels, Ensemblex uses the cells with a consensus assignment across the three remaining tools (e.g., Demuxlet, Souporcell, and Vireo-GT) as a proxy for ground-truth. The balanced accuracy for each tool is calculated using equation 1:

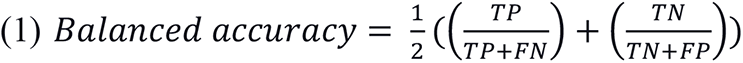

Where TP is the number of correctly classified singlets; true-negative (TN) is the number of correctly classified doublets; FP is the number of incorrectly classified singlets; false-negative (FN) is the number of incorrectly classified doublets. The probability distribution of each constituent tool (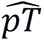) is then weighted by its estimated balanced accuracy (*w*_*j*_) to produce an accuracy-weighted ensemble probability for each cell:

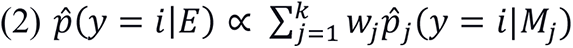

Where 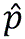 is the probability that a barcode belongs to class *i*; *y* is the class variable with *c* possible values, *y* ∈ (1, …, *c*); *c* is the number of pooled samples plus 1 to account for doublets; *E* is a vector of the results of *M* classifiers, *E* = (*M*_1_, …, *M*_*k*_); *M*is the individual constituent demultiplexing output from each tool. Given 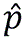, Ensemblex assigns each barcode’s sample identity (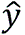) as the class (sample label) with the maximum probability:

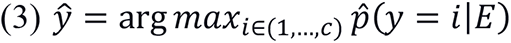

### Step 2: Graph-based doublet detection

Ensemblex employs a graph-based approach to identify doublets that are incorrectly labeled as singlets by the accuracy-weighted probabilistic ensemble component (Step 1). For graph-based doublet detection, Ensemblex leverages pre-defined features returned from each constituent tool:

1. Demuxalot: doublet probability;
2. Demuxlet/Freemuxlet: singlet log likelihood – doublet log likelihood;
3. Demuxlet/Freemuxlet: number of single nucleotide polymorphisms (SNP) per cell;
4. Demuxlet/Freemuxlet: number of reads per cell;
5. Souporcell: doublet log probability;
6. Vireo: doublet probability;
7. Vireo: doublet log likelihood ratio.

For each feature independently, the pooled cells are ordered from the most to the least probable doublet and are assigned a percentile rank. Beginning with a percentile threshold of 99.99, Ensemblex screens each cell to identify those that exceed the percentile threshold across all features; cells that exceed the percentile threshold across all features are labeled as “confident doublets”. For each iteration, Ensemblex decreases the percentile threshold by 0.01 and repeats the screening process until it has identified *n* confident doublets (nCD). Ensemblex performs a parameter sweep to determine the optimal nCD to use for graph-based doublet detection (see below).

Next, the above features are input into a PCA using the *stats* (v3.6.2) R package (25) and a Euclidean distance matrix is generated from the first two principal components (PC). For each confident doublet independently, the remaining cells in the pool are assigned a percentile rank based on their proximity in Euclidean space to the confident doublet and the cells that exceed the designated nearest neighbour percentile threshold (pT) are identified. For all cells that exceeded the designated pT for any confident doublet (putative doublets), Ensemblex computes the number of times the putative doublet was amongst the nearest neighbours of any confident doublet (fNN); an fNN equal to nCD indicates that a putative doublet was amongst the top nearest neighbours for each confident doublet.

To optimize the nCD and pT parameters for experimentally pooled samples lacking ground-truth labels, Ensemblex performs an automated parameter sweep at each pairwise combination of nCD and pT values; nCD values range from 50 to 300, in increments of 50, while pT values depend on the expected doublet rate (exDR) and range from 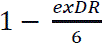 to 1 − *exDR*, in intervals of 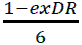. The distribution of fNN values for each combination of nCD and pT parameters are plotted and Pearson’s measure of kurtosis (K), is used to predict which combination of pT and nCD values optimize the identification of true doublets while minimizing the rate of incorrectly labelled true singlets as doublets. Ensemblex screens for combinations of nCD and pT values that result in negatively skewed fNN distributions with high K, signifying high peakedness and heavy tails. High peakedness indicates that cells exceeding the designated pT concentrated around nCD, reflecting their proximity in Euclidean space to all high confident doublets, while heavy tails indicate that even cells with lower fNN values were identified as nearest neighbour to many confident doublets. Ensemblex first identifies the pT that returns the highest K, on average, across nCD values tested in the parameter sweep using equation 4:

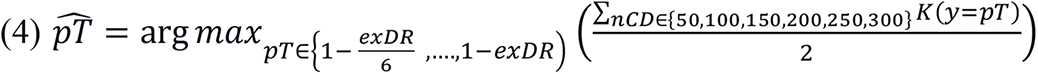

Where K of the distribution of fNN values of the putative doublets is defined as:

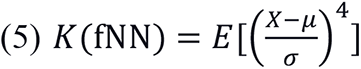

Where μ is the mean of the distribution and σ is the standard deviation. Upon identifying the optimal pT value (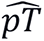), Ensemblex plots the K corresponding to 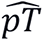 across all nCD values tested in the parameter sweep. If an inflection point is identifiable, Ensemblex identifies 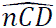 as the nCD value corresponding to the point of inflection on the curve. Otherwise, Ensemblex identifies 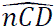 as the nCD value corresponding to the highest K. Cells flagged as putative doublets identified using 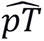 and 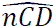 are labelled as doublets by Ensemblex.

### Step 3: Ensemble-independent doublet detection

Benchmarking on computationally multiplexed pools with known ground-truth sample labels revealed that certain genetic demultiplexing tools, namely Demuxalot and Vireo, showed high doublet detection specificity, but that Steps 1 and 2 of the Ensemblex workflow failed to correctly label a subset of doublet calls by these tools. To mitigate this issue and maximize the rate of doublet identification, Ensemblex labels the cells that are identified as doublets by Vireo or Demuxalot as doublets by default; however, users can nominate different tools for the ensemble-independent doublet detection component depending on the desired doublet detection stringency. Doublet specificity was computed using equation 6:

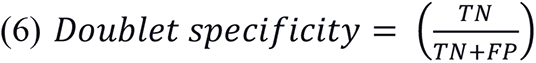

Where TN is the number of correctly classified doublets; FP is the number of true singlets incorrectly classified as doublets.

### Ensemblex singlet assignment confidence score

Ensemblex computes a singlet confidence score to inform which cells should be discarded to avoid misclassification in downstream analyses. First, Ensemblex evaluates how well an individual constituent tool’s assignment probability (e.g., Demuxalot) corresponded to the accuracy of their assignment, using consensus cells across the three remaining tools (e.g., Demuxlet, Souporcell, Vireo) as a proxy for ground-truth, by fitting a binary logistic regression model to compute the odds that a singlet was correctly classified given its corresponding probability. Using the binary logistic regression models, Ensemblex computes the AUC using the empirical method implemented in the *ROCit* (v2.1.1) R package for each tool (26). Then, for each cell, if Ensemblex’s sample label matches that of a constituent tool, and if the assignment probability of the constituent tool supersedes its probability threshold, the tool’s computed AUC is added to the accuracy-weighted probabilistic ensemble probability produced in Step 1 to yield the confidence score. By default, singlet assignments with a confidence score less than 1.00 are labelled as unassigned by Ensemblex. Ensemblex’s confidence score and the designated threshold is a successful predictor of accurately classified singlets because singlets will only achieve a confidence score ≥ 1 if:

1. All constituent tools show the same sample label (accuracy-weighted probabilistic ensemble probability = 1.00);
2. At least one constituent tool confidently assigns the cell to an individual donor and the constituent tool’s probability assignment adequately corresponds to the overall accuracy of their singlet assignment.

### Application of Ensemblex with and without prior genotype information

Given the dependencies of certain tools on prior genotype information, there are notable differences between the Ensemblex workflows for demultiplexing with and without prior genotype information. When demultiplexing with prior genotype information, Ensemblex leverages the sample labels from Demuxalot, Demuxlet, and Vireo-GT with prior genotype information, and Souporcell without prior genotype information. When demultiplexing without prior genotype information, Ensemblex leverages the sample labels from Demuxalot, Freemuxlet, Souporcell, and Vireo. However, given that Demuxalot requires prior genotype information, Ensemblex uses the estimated donor .vcf file generated by Freemuxlet for input into the Demuxalot algorithm as prior genetic data.

### Running the Ensemblex pipeline

A complete user guide for running the Ensemblex pipeline can be found at the Ensemblex GitHub site: https://neurobioinfo.github.io/ensemblex/site/. We provide two distinct yet highly comparable pipelines depending on the availability of prior genotype information. Both pipelines can be downloaded as a singularity image and are comprised of four steps:

1. Establish the pipeline and working directory;
2. Prepare input files for constituent genetic demultiplexing tools;
3. Parallel demultiplexing by constituent genetic demultiplexing tools;
4. Application of the Ensemblex algorithm for ensemble classification.

As input into the Ensemblex pipeline, users must provide a .tsv file describing the barcodes of the pooled cells, a. bam sequencing file for the pool, a reference genotype .vcf file (e.g., 1000 Genome Project) (27), a reference genome sequence .fasta file (e.g., 10X Genomics), and, if demultiplexing with prior genotype information, a .vcf file describing the genetic data of the pooled samples.

### Genetic demultiplexing by constituent tools

Genetic demultiplexing by the constituent demultiplexing tools was performed following best practices as defined by the authors of the respective tools using Python (v3.8.10).

### Demuxalot

CellRanger-generated .bam file, filtered barcode .tsv file, and the corresponding donor .vcf file were used as input into the Demuxalot workflow. Candidate variants for scRNAseq genotyping were retained if the minimum coverage was > 200 and minimum alternative coverage was > 10. The top 100 SNPs per donor were retained to cluster the cells by genotype. Doublet calls were made with a prior strength of 0.25.

### Demuxlet

We used the popscle suite (https://github.com/statgen/popscle) for Demuxlet. CellRanger-generated .bam file, filtered barcode .tsv file, and the corresponding donor .vcf file were used as input into the Demuxlet workflow. The *dsc-pileup* function was first used to pileup candidate variants around known variant sites with the following parameters: ––cp-BQ 40 ––min-BQ 13 ––min-MQ 20 ––minTD 0 ––min-total 0 ––min-uniq 0 ––min-snp 0. The Demuxlet algorithm was then applied to cluster the cells by genotype with the following parameters: ––geno-error-offset 0.10 ––geno-error-coeff 0.00 ––min-callrate 0.50 ––doublet-prior 0.50 ––cap-BQ 40 ––min-BQ 13 ––min-MQ 20 ––min-TD 0 ––min-total 0 ––min-uniq 0 ––min-snp 0.

### Freemuxlet

We used the popscle suite (https://github.com/statgen/popscle) for Freemuxlet. CellRanger-generated .bam file, filtered barcode .tsv file, and reference genotype .vcf file from the 1000 Genomes Project, phase 3 (27), were used as input into the Freemuxlet workflow. The *dsc-pileup* function was first used to pileup candidate variants around known variant sites with the following parameters: ––cp-BQ 40 ––min-BQ 13 ––min-MQ 20 ––minTD 0 ––min-total 0 ––min-uniq 0 ––min-snp 0. The Freemuxlet algorithm was then applied to cluster the cells by genotype with the following parameters: ––doublet-prior 0.50 ––bf-thres 5.41 ––frac-init-clust 0.50 ––inter-init 10 ––cap-BQ 40 ––min-BQ 13 ––min-total 0 ––min-uniq 0 ––min-snp 0.

### Souporcell

CellRanger-generated .bam file, filtered barcode .tsv file, 10X Genomics reference .fasta file, and the corresponding donor .vcf file when demultiplexing with prior genotype information were used as input into the Souporcell workflow. A FASTQ file was first generated from the .bam file using the *renamer.py* script. These reads were mapped to the reference genome using minimap2 with the following parameters: ––ax splice –t 8 –G50k –k 21 –w 11 –sr ––A2 –B8 – O12,32 –E2,1 –r200 –p.5 –N20 –f1000,5000 –n2 –m20 –s40 –g200 –2k50m –secondary=no. The barcodes and UMI were added back to the .sam file using the *retag.py* script and the resulting .bam file was sorted and indexed with Samtools. Variants were called using Freebayes with the following parameters: ––iXu –C 2 –q 20 –n 3 –E 1 –m 30 –min-coverage 6. Vartix was used to compute the number of alleles for each cell using the following parameters: ––umi – mapq 30 –scoring-method coverage. The Souporcell algorithm was then applied to cluster the cells by genotype; when demultiplexing with prior genotype information the –– known_genotypes and ––known_genotypes_sample_names parameters were included. Troublet was used to identify doublets and the *consensus.py* script was used for genotype and ambient RNA co-inference.

### Vireo

CellRanger-generated .bam file, filtered barcode .tsv file, reference genotypes from the 1000 Genomes Project, phase 3 (27), and the corresponding donor .vcf file when demultiplexing with prior genotype information were used as input to the Vireo workflow. CellSNP was used to identify candidate variants for scRNAseq genotyping with the following parameters: –– minMAF 0.1 and ––minCOUNT 100. The Vireo algorithm was then applied to cluster the cells by genotype with the ––forceLearnGT parameter; when demultiplexing with prior genotype information (Vireo-GT) the ––d and ––t GT parameters were used.

### Consensus demultiplexing framework

For the consensus demultiplexing framework, singlets were considered confidently classified if Demuxalot, Demuxlet, Vireo, and Souporcell assigned a cell to the same donor-of-origin. Cells classified as “ambiguous” or doublet by at least one tool were discarded.

### Generation of computationally pooled samples for ground-truth benchmarking

To benchmark Ensemblex on computationally pooled samples with known ground-truth sample labels, we leveraged 80 independently sequenced iPSC lines from Parkinson’s disease patients and healthy controls, which were differentiated towards a dopaminergic neuronal state and sequenced after 65 days of differentiation as part of the FOUNDIN-PD (14). Controlled access FASTQ files from the independently sequenced iPSC lines were obtained from https://www.ppmi-info.org/ (accessed 09-17-2023) and processed by the CellRanger *counts* pipeline (v3.1.0) with default parameters and aligned to GRCh38 reference genome. The CellRanger-generated .bam and filtered barcode files were used as input into the *synth_pool.py* script produced by the authors of Vireo to simulate sample pooling (9). In brief, reads from a subset of cells from the iPSC line-specific .bam files were merged and doublets were generated by combining the reads from random cell pairs. Sample identities were added to each cell’s barcode, revealing the ground-truth sample labels for benchmarking procedures.

To evaluate how genetic demultiplexing performance varied as a function of the number of multiplexed samples, we generated 96 computationally multiplexed pools using the 80 FOUNDIN-PD lines with sample sizes of 4, 8, 16, 24, 32, 40, 48, 56, 64, 72, and 80. An equal number of cells from each line were used in the *in silico* pool. For the sample size of four we generated six replicates; for the sample sizes of 8-80 we generated nine replicates each. Replicates were produced with different sample and cell combinations. The 96 *in silico* pools averaged 17,396 cells (minimum = 8,696; maximum = 26,087). For this experiment, we maintained a 15% doublet rate as previously described (9).

To evaluate how genetic demultiplexing performance varied as a function of the number of cells in a pool, we generated 18 computationally multiplexed pools using the 80 FOUNDIN-PD lines with 8,000, 16,000, 24,000, 32,000, 40,000, and 48,0000 pooled cells; we generated three replicates per pool size. Twenty-four samples were multiplexed for each pool and an equal number of cells from each sample were used. Replicates were produced with different sample and cell combinations. For this experiment, we simulated a doublet rate of 6% per 8,000 pooled cells.

To evaluate if the overall demultiplexing performance varied due to the underrepresentation of a cell line, we generated 15 computationally multiplexed pools using the 80 FOUNDIN-PD lines comprising 23 multiplexed samples with 1,000 cells and one randomly selected sample that showed various degrees of underrepresentation, including 100 cells (10%), 300 cells (30%), 500 cells (50%), 700 cells (70%), or 900 cells (90%). Three replicates were generated for each degree of underrepresentation. Replicates were produced with different sample and cell combinations. For this experiment, we maintained a 18% doublet rate.

WGS for the 80 donors from which the FOUNDIN-PD lines were derived was performed on whole blood-extracted DNA as previously described by the Parkinson’s Progression Markers Initiative (PPMI) (28). The controlled-access WGS .vcf files were obtained from https://www.ppmi-info.org/ (accessed 09-17-2023). Genotypes of common variants (minor allele frequency > 5%) were used as prior genotype information for the genetic demultiplexing tools in the benchmarking analyses.

### Preparation, processing, and analysis of experimentally pooled samples

Unless specified otherwise, experimentally pooled samples were processed with the CellRanger *counts* pipeline (v5.0.1) and analyzed with the *Seurat* (v5.0.0) R package (29), using the scRNAbox analytical pipeline (30).

### Non-small cell lung cancer dataset

NSCLC dissociated tumor cells from seven donors were labelled with TotalSeq-B Human TBNK Cocktail (18). Multiplexed cells were then sequenced on an Illumina NovaSeq 6000 to an average read depth of approximately 70,000 reads per cell for gene expression and 25,000 reads per cell for CellPlex. Publicly available gene expression .bam and barcode .tsv files returned from the CellRanger *multi* pipeline (v6.1.2) were obtained from the 10X Genomics Datasets portal (<u>10X Genomics Datasets</u>) and used as input into the Ensemblex pipeline. We used the sample-specific gene expression .bam files and the BCFtools (v1.16) *mpielup* function to generate genotype likelihoods for prior genotype information (31).

We used HTOdemux to assign the cells back to their donor-of-origin based on the CMO expression profiles as a proxy for ground-truth sample labels (19). Publicly available feature-barcode expression matrices were filtered to only include CMO labels used for multiplexing — CMO301, CMO302, CMO303, CMO304, CMO306, CMO307, and CMO308 — and barcodes with a CMO count > 0. The CMO expression profiles were normalized with Seurat’s *NormalizeData* function using the CLR normalization method and HTOdemux was applied to the CMO assay using a positive quantile of 0.99.

### Dopaminergic neuron dataset

Jerber et al. sequenced multiplexed experiments comprising 22 healthy donor iPSC lines from the HipSci project (32) (http://www.hipsci.org) on days 11, 30, and 52 of DaN differentiation using Illumina HiSeq 4000 to an average depth of 40,000-60,000 reads per cell (12). We used three technical replicates for each timepoint, which are comprehensively described in **Additional File 1: Table S3**. Publicly available gene expression .fastq files were obtained from the European Nucleotide Archive (ENA) with accession number ERP121676 and processed with the CellRanger *counts* pipeline (v5.0.1) with default parameters using the GRCh37 reference genome. The CellRanger-generated. bam files, filtered barcode .tsv files, and .vcf files describing the pooled samples (see below) were used as input into the Ensemblex pipeline for each technical replicate independently. Filtering of the scRNAseq data was performed as described by Jerber et al. (12). Genes with non-zero counts in at least 0.05% of cells were retained. DoubletFinder (v2.0.4) was applied independently to each technical replicate. Time-point specific replicates were integrated with Seurat’s integration algorithm (33) and clustered by the Louvain network detection using the top 50 PCs and 10 nearest neighbours.

Whole-exome sequencing (WES) .vcf files corresponding to the 22 pooled HipSci lines were obtained from the ENA with accession number PRJEB7243 (34). Genotypes of common variants (minor allele frequency > 1%) were used as prior genotype information for the genetic demultiplexing tools (12).

### Neural stem cell dataset

We performed two multiplexed experiments comprising iPSCs from individuals with ADHD and heathy controls differentiated into NSCs: Experiment 1 (*n* ADHD = 7; *n* control = 6) and Experiment 2 (*n* ADHD = 9; *n* control = 7).

### Subject recruitment

Patients diagnosed with ADHD and matching healthy controls between 6−18 years old were recruited by the Department of Child and Adolescent Psychiatry and Psychotherapy of the University of Zurich, as described previously (35). Inclusion and exclusion criteria for recruitment of these individuals described previously (35). **Additional File 1: Table S4** provides a list of the individual subjects and their derived cell lines included in this study. Salivary DNA from ADHD patients and controls was genotyped using the Infinium Global Screening Array (Illumina), as previously described, and used as prior genotype information for genetic demultiplexing (35).

### Neural stem cell culture

The generation and characterization of iPSC used in this study and the NSCs differentiation protocols were previously described in (35) (36). NSCs cultures were seeded in two independent experiments (designated as “1” and “2”), each of them consisting of NSCs pooled together into two culture dishes and maintained as NSCs until 100% confluence, when all iPSC lines were combined into one sample for sequencing. For most cell lines different clones for each iPSC line were used in the two experiments **Additional File 1: Table S5**. When applicable, the second clones of the same NSCs lines were cultured separately (designated as “.1” and “.2”) in a second experiment. In the first experiment, 56,250 cells per cell line were seeded in the pooled dishes. In the second experiment the proportions of cells seeded we adjusted to their proliferation profile assessed in (36). Upon reaching 100% confluence, cells were dissociated for scRNAseq experiments and combined to a single sample for sequencing as described below.

### Dissociation of pooled neural stem cell cultures for single-cell RNA sequencing

Cells were washed in PBS and then incubated with 1 mL of StemPro Accutase (Gibco) for 3 minutes at 37°C. After incubation, 2 mL of PBS, stopping the Accutase reaction, and cells were gently pipetted up and down between 5 to 10 times to break up clumps of cells before transfer to a 15 mL conical tube. The cells were centrifuged at 300 x g for 5 minutes and the supernatant was removed. Following, 334 µL of Neural Expansion Media (NEM) was added to each cell pellet using a 1000 µL pipette tip until cells were completely resuspended. An additional 666 µL of NEM was added to each well and gently pipette mixed 5 times. A 100-µm cell strainer was used to filter the cell suspension before centrifugation at 300 x g for 4 minutes. The supernatant was carefully removed, and the pellet was resuspended in 3 mL of PBS 1x containing 0.04% Bovine Serum Albumin (BSA) by pipetting up and down 5 times using a 5 mL serological pipette. The cells were centrifuged at 300 x g for 10 minutes and further submitted to live cell sorting with the Magnetic Dead Cell Removal Kit (Miltenyi Biotec, 130-090-101), according to the manufacturer. The resulting flow-through containing live cells was centrifuged for 300 x g for 5 minutes and the supernatant was removed carefully to not disturb the cell pellet. Cells were resuspended in 1 mL of PBS 1x containing 0.04% BSA for automated cell counting. For each experiment, the cells from the two culture dishes were processed in parallel. Equal counts of cells were combined for the final cell suspension for scRNAseq preparation at the Functional Genomics Center Zurich at the University of Zurich.

### Library processing and sequencing

All samples were processed using the 10x Genomics Chromium 3’ Single Cell Protocol and sequenced using NovaSeq 6000 S1 (Illumina). For the first sample containing NSC pools 1.1 and 1.2, 18,000 NSCs were loaded into one single 10x Genomics Lane to target 13,000 cells. For the second sample containing NSC pools 2.1 and 2.2, 29,000 NSCs were loaded to target 18,000 cells.

### Demultiplexing and scRNAseq analysis

FASTQ files were processed with the CellRanger *counts* pipeline (v5.0.1) with default parameters and aligned to the GRCh37 reference genome. The CellRanger-generated. bam files, filtered barcode .tsv files, and .vcf files describing the pooled samples were used as input into the Ensemblex pipeline. Genotypes of common variants (minor allele frequency > 1%) were used as prior genotype information for the genetic demultiplexing tools. The filtered feature-barcode expression matrices were used to analyze the pooled cells following a standard scRNAseq analysis workflow using Seurat (30). Cells were filtered for > 500 total and unique RNA transcripts. Doublets were removed using DoubletFinder (v2.0.4). The two NSC samples were integrated using Seurat’s integration algorithm (33). The top 25 PCs were selected for Louvain network detection to identify clusters using 65 nearest neighbours. Twelve clusters were identified at a clustering resolution of 0.25, which were assigned as eight putative cell types using a combination of known markers and gene enrichment analysis. The top marker genes from each cluster were identified using Seurat’s *FindAllMarkers* with the Wilcoxon rank-sum test. Significant DEGs (log2 fold change > 0.25 and P-value < 0.05) were input into EnrichR (37) and cell types were predicted with the *Cell Marker Augmented 2021* (38) and *Azimuth Cell Types 2021* (39) libraries. Multiple clusters showed expression profiles for similar broad cell types — Neurons, NPCs, and NSCs. We used Seurat’s *FindMarkers* function to identify differentially expressed marker genes between the clusters of the same broad cell type and top marker genes were selected to identify the cell subtypes.

For each putative cell type, DGE was calculated between ADHD and controls using the MAST statistical framework (22, 40). Pooled cells were assigned as ADHD or control based on the demultiplexed sample labels from each of the individual genetic demultiplexing tools. Cells labeled as “ambiguous singlets” or doublets by the individual tools were excluded from their respective DGE analysis. P-values were corrected for multiple hypothesis testing using the Bonferroni method. A gene was considered differentially expressed if the adjusted P-value was ≤ 0.01 and the absolute value of the Log2 fold-change was ≥ 0.5. To compute DGE using the sample labels from the individual tools after the removal of Ensemblex’s putative doublet calls, we repeated the above procedures but this time all cells labeled as doublets by the respective tool or Ensemblex were excluded from the DGE analysis.

### Performance metrics and statistical analyses

We performed all statistical analyses using the R statistical software (v4.2.2) (41). We used the *ggplot2* R package (v3.4.2) for data visualization (42).

### Singlet classification

A singlet was considered correctly classified if the demultiplexed sample label matched the ground-truth sample label (i.e., specific sample ID) and the assignment probability exceeded the recommended threshold for the respective tool. For computationally multiplexed pools, the proportion of correctly classified singlets was computed as:

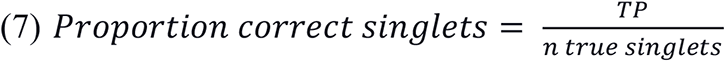

For the NSCLC dataset, HTOdemux’s sample labels were considered ground-truth, and the singlet TP and FP rate were computed as:

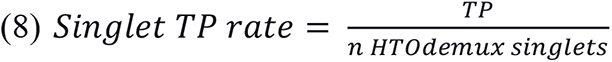

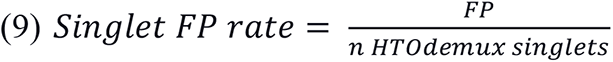

### Doublet identification

A doublet was considered correctly classified if the demultiplexed sample label matched the ground-truth sample label, independent of the assignment probability. For computationally multiplexed pools, the proportion of correctly classified doublets was computed as:

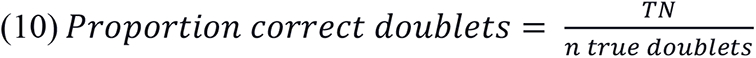

For the NSCLC dataset, TP doublets were defined as cells classified as doublets by both HTOdemux and Ensemblex; FP doublets were defined as cells classified as singlets by HTOdemux and doublets by Ensemblex; FN doublets were defined as cells classified as doublets by HTOdemux and singlets by Ensemblex. The doublet TP, FP, and FN rates were computed as:

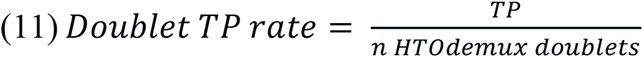

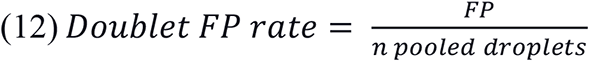

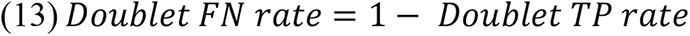

### Adjusted Rand Index

To evaluate the similarity between two distinct sample clusterings we computed the ARI using the *pdfCluster* (v1.0.4) R package (43). For the benchmarking analyses, we computed the ARI between the demultiplexed sample labels by each genetic demultiplexing tool and the ground-truth sample labels (computationally pooled samples) or HTOdemux’s sample labels (NSCLC dataset). We followed the same procedure when computing the ARI between Ensemblex’s sample labels and those of its constituent tools (DaN and NSC datasets); however, the ground-truth sample labels were replaced by Ensemblex’s sample labels for these analyses. For experiments evaluating the impact of doublets on the stability of clusters in gene expression space, we computed the ARI between clusters at a given clustering resolution after removing doublets identified by each genetic demultiplexing tool. Clustering stability was computed at resolutions of 0.05, 0.1, 0.2, 0.3, 0.4, 0.5, 0.6, 0.7, 0.8, 0.9, and 1.0. For each clustering resolution, 25 iterations of Louvain clustering were performed while shuffling the order of the nodes in the graph. The ARI between clustering pairs at each clustering resolution was then computed.

### Balanced accuracy

Balanced accuracies were computed to evaluate the binary classification performance of each genetic demultiplexing tool on imbalanced datasets, where doublets represented a minority class compared to singlets. The balanced accuracy of each genetic demultiplexing tool was computed against the ground-truth sample labels (computationally pooled samples) or HTOdemux’s sample labels (NSCLC dataset) using equation 1.

### Matthew’s correlation coefficient (MCC)

The MCC was used as a second metric for evaluating the binary classification performance of the genetic demultiplexing tool. The MCC of each genetic demultiplexing tool was computed against the ground-truth sample labels (computationally pooled samples) or HTOdemux’s sample labels (NSCLC dataset) using equation 14:

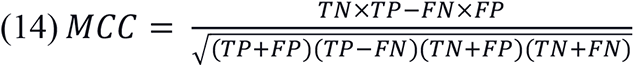

### Area under the receiver operating characteristic curve for singlet detection

To evaluate how well each genetic demultiplexing tool’s assignment probability corresponded to the accuracy of their singlet assignments when ground-truth sample labels were known, we fit a binary logistic regression model to compute the odds that a singlet was correctly classified by a tool given the corresponding confidence score or probability. Correctly and incorrectly classified singlets were set as the positive and negative references, respectively. We then used the binary logistic regression model to compute the receiver operating characteristic curve for each tool, which plots the singlet TP and FP rates across classification thresholds, and calculated the AUC using the empirical method implemented in the *ROCit* (v2.1.1) R package (26).

## Declarations

### Ethics approval and consent to participate

The iPSC lines (ADHD & controls) used in this project were approved by the Cantonal Ethics Committee Zurich (BASEC-Nr.-2016-00101 & BASEC-Nr.-201700825) and followed the latest version of the Declaration of Helsinki, as previously reported (35). The subjects and/or parents have voluntarily consented to participate in this study.

### Consent for publication

Not applicable.

### Availability of data and materials

Transcriptional data for the 80 independently sequenced iPSC lines and the corresponding WGS data are available from the PPMI database (www.ppmi-info.org/access-dataspecimens/download-data), RRID:SCR 006431. For up-to-date information on the study, visit www.ppmi-info.org. Processed transcriptional data for the NSCLC dataset are available from the 10X Genomics Datasets Portal (https://www.10xgenomics.com/datasets/20k-mixture-of-nsclc-dtcs-from-7-donors-3-v3-1-with-intronic-reads-3-1-standard). Transcriptional data for the DaN datasets are available from the ENA with accession number ERP121676. WES data for the 22 HipSci lines pooled in the DaN datasets are available from the ENA with accession number PRJEB7243. Processed scRNAseq data for the NSC dataset are available from the corresponding author upon reasonable request. The code used for the analyses presented in the work are available at https://github.com/neurobioinfo/ensemblex. Ensemblex is freely available under an MIT open-source license at https://zenodo.org/records/11639103.

## Authors’ contributions

MRF, EAF, RAT, and SMKF conceived the study. MRF developed the Ensemblex framework and wrote the corresponding R code. MRF performed the analyses and produced the figures. MRF and SA developed the Ensemblex pipeline and created the GitHub site. MRF and SA tested the Ensemblex pipeline. MRF wrote the Ensemblex documentation. MRF, AAD, RAT and SMKF interpreted the datasets. CMYO performed all cell cultures and sequencing preparation for the NSC dataset. MRF, CMYO, and RAT performed the cell type annotations for the NSC dataset. LS and EG provided the NSC genetic data. SW recruited the subjects for the NSC dataset. MRF wrote the manuscript with input from all authors. EG supervised the NSC data collection. RAT and SMKF supervised the project.

## Abbreviations

ADHD: attention deficit hyperactivity disorder
ANOVA: Analysis of variance
ARI: Adjusted Rand Index
AUC: area under the receiver operating characteristic curve
BSA: Bovine Serum Albumin
CMO: Cell Multiplexing Oligonucleotides
DaN: dopaminergic neurons
DGE: differential gene expression
DEG: differentially expressed genes
ENA: European Nucleotide Archive
eQTL: expression quantitative trait loci
FN: false-negative
fNN: nearest neighbour frequency
FOUNDIN-PD: Foundational Data Initiative for Parkinson’s Disease
FP: false positive
iPSC: induced pluripotent stem cell
K: kurtosis
MAST: model-based analysis of single-cell transcriptomics
MCC: Matthew’s Correlation Coefficient
nCD: number of confident doublets
NEM: neural expansion media
NPC: neural progenitor cell
NSC: neural stem cell
NSCLC: non-small cell lung cancer
PC: principal component
PCA: principal component analysis
PPMI: Parkinson’s Progression Markers Initiative
pT: nearest neighbour percentile threshold
scATACseq: single-cell assay for transposase-accessible chromatin sequencing
scRNAseq: single-cell RNA sequencing
SNP: single nucleotide polymorphism
snRNAseq: single-nuclei RNA sequencing
TN: true-negative
TP: true-positive
UMI: unique molecular identified
WES: whole-exome sequencing
WGS: whole-genome sequencing

## Acknowledgments

The author’s acknowledge Dan Spiegelman for his help in processing the VCF files for individuals pooled in the NSC dataset. Schematic illustrations presented in the manuscript were prepared with BioRender (https://www.biorender.com/).

## Authors’ information

This work was supported by the Michael J Fox Foundation in a grant award to EAF, RAT, and SMKF [MJFF-021629]. MRF is supported by a CIHR Canada Graduate Scholarships-Master’s Award, a Fonds de Recherche Santé Québec Master’s Award, and a Brain Canada Rising Stars Award. EAF is supported by a Fonds d’Accéleration des Collaborations en Santé (FACS) grant from CQDM/MEI and a Canada Research Chair (Tier 1) in Parkinson’s disease. R.A.T. received funding through the McGill Healthy Brains for Healthy Lives (HBHL) Postdoctoral Fellowship and Molson NeuroEngineering Fellowship. SMKF received funding from Brain Canada and the Montreal Neurological Institute-Hospital. CMYO is supported by the Candoc PhD grant from the University of Zurich (UZH) [FK-22-044].

