## Additional File 1 for "Ensemblex: an accuracy-weighted ensemble genetic demultiplexing framework for population-scale scRNAseq sample pooling"

### Supplemental Figures and Tables

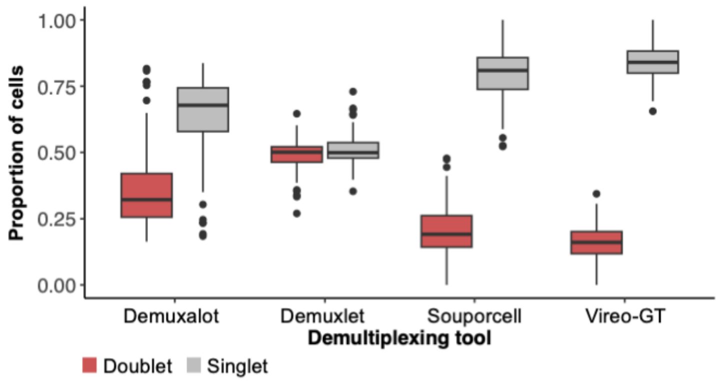


**Figure S1. Proportion of doublets and singlets classified correctly by the indicated tool amongst cells correctly classified by only one tool.** Genetic demultiplexing tools using prior genotype information were evaluated on 96 *in silico* pools with known ground-truth sample labels ranging in size from 4 to 80 multiplexed induced pluripotent stem cell (iPSC) lines from genetically distinct individuals, averaging 17,396 cells per pool and a 15% doublet rate.

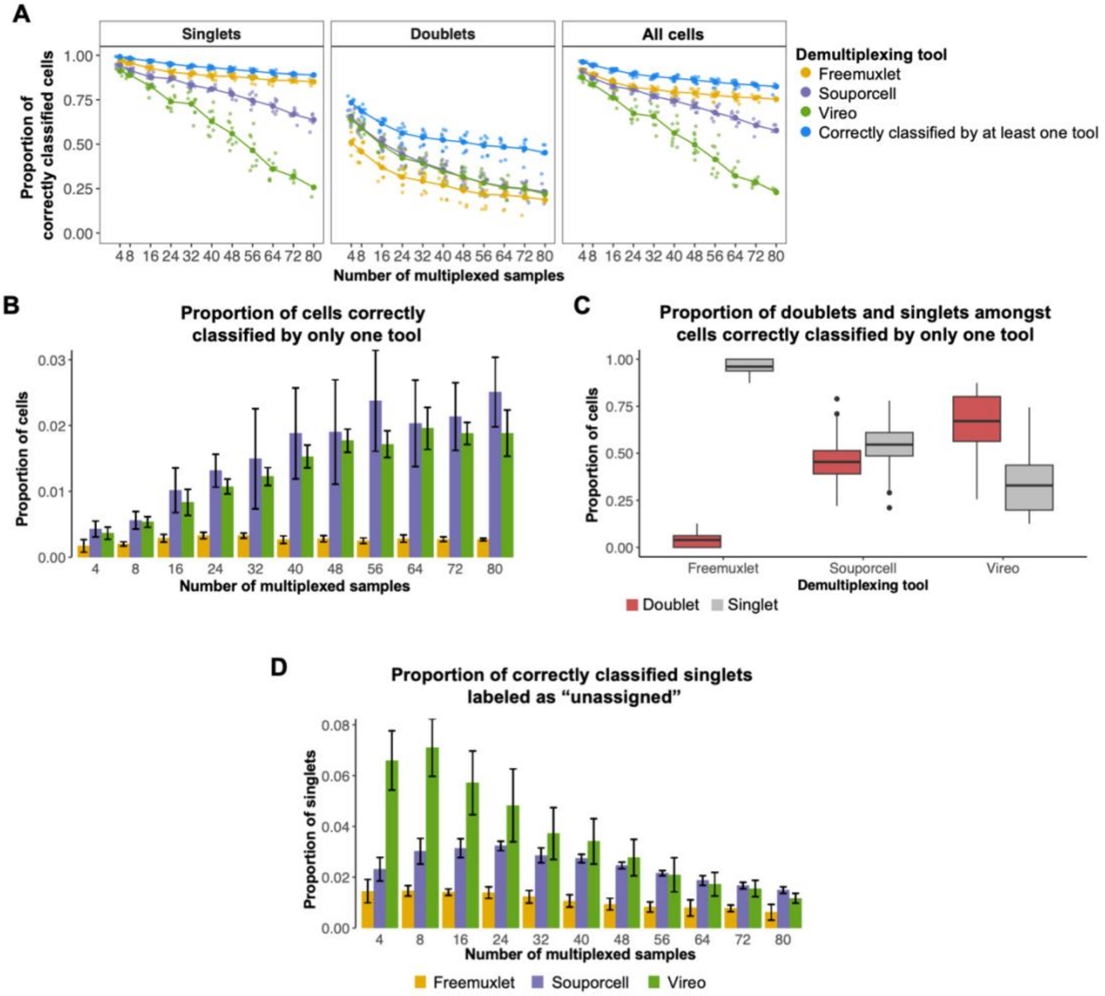


**Figure S2. Evaluation of existing individual genetic demultiplexing tools when demultiplexing without prior genotype information.** Evaluation of genetic demultiplexing tools with prior genotype information on 96 *in silico* pools with known ground-truth sample labels ranging in size from 4 to 80 multiplexed induced pluripotent stem cell (iPSC) lines from genetically distinct individuals, averaging 17,396 cells per pool and a 15% doublet rate. **A)** Line graphs showing the proportion of correctly classified singlets, doublets, and all cells by each individual genetic demultiplexing tool across varying numbers of multiplexed iPSC lines in a single pool (sample number). The large dots show the mean proportion of correct classifications by an individual tool across replicates at a given sample size (n = 9 per pool size). The blue points show the proportion of cells that were correctly classified by at least one individual genetic demultiplexing tool: Freemuxlet, Souporcell, or Vireo. **B)** Bar chart showing the mean proportion of total cells from an individual pool correctly classified by only one genetic demultiplexing tool. Error bars represent one standard deviation from the mean. (n = 9 per pool size) **C)** Boxplots showing the proportion of doublets and singlets classified correctly by the indicated tool amongst cells correctly classified by only one tool (n = 96 pools). **D)** Bar chart showing the proportion of correctly classified singlet cells labelled as "unassigned" (ambiguous singlet assignments) due to assignment probabilities below the recommended threshold of the respective genetic demultiplexing tool. Error bars represent one standard deviation from the mean. (n = 9 per pool size).

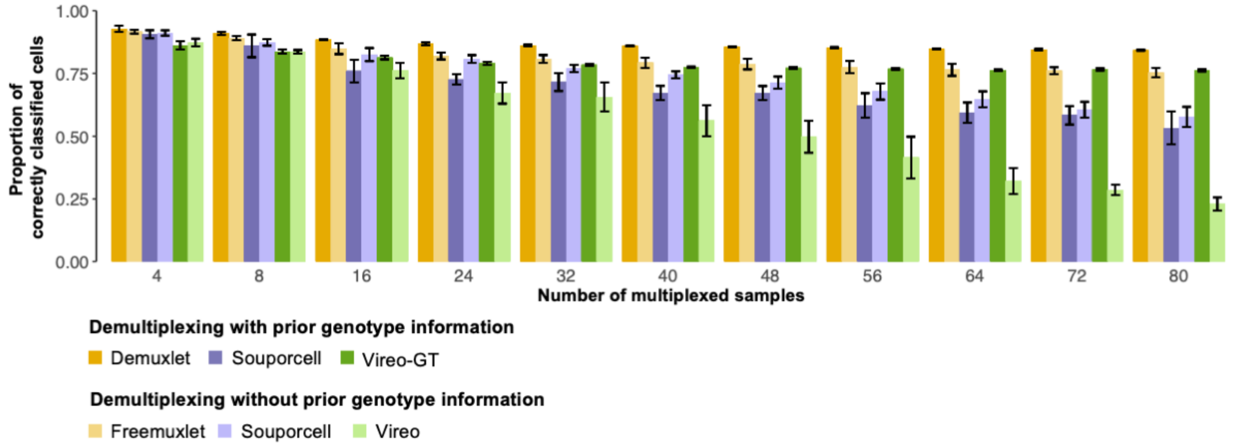


**Figure S3. Comparison of genetic demultiplexing performance when demultiplexing with and without prior genotype information.** Genetic demultiplexing tools were evaluated 96 *in silico* pools with known ground-truth sample labels ranging in size from 4 to 80 multiplexed induced pluripotent stem cell (iPSC) lines from genetically distinct individuals, averaging 17,396 cells per pool and a 15% doublet rate. Error bars represent one standard deviation from the mean.

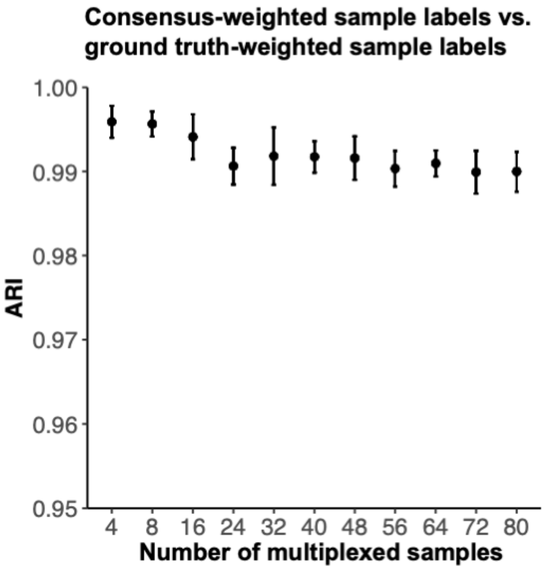


**Figure S4. Validating the accuracy-weighted probabilistic ensemble component of the Ensemblex framework on *in silico* pools with known ground-truth sample labels.** The accuracy-weighted probabilistic ensemble component (Step 1) of the Ensemblex framework weights the assignment probabilities from the constituent genetic demultiplexing tools by their estimated balanced accuracy for the dataset and the weighted probabilities are used to determine the most probable sample labels for each cell. The estimated balanced accuracy of an individual tool (e.g., Demuxalot) is computed on cells with a consensus sample label across the three remaining tools (e.g., Demuxlet, Souporcell, and Vireo), which Ensemblex uses as a proxy for ground-truth sample labels. To validate our approach of using the balanced accuracy derived from consensus cells for weighting each tool's assignment probabilities, we leveraged 96 *in silico* pools with known ground-truth sample labels ranging in size from 4 to 80 multiplexed induced pluripotent stem cell (iPSC) lines from genetically distinct individuals, averaging 17,396 cells per pool and a 15% doublet rate. The figure shows the mean Adjusted Rand Index (ARI) between consensus-weighted sample labels and ground-truth weighted sample labels across replicates at a given sample size. Error bars represent one standard deviation from the mean.

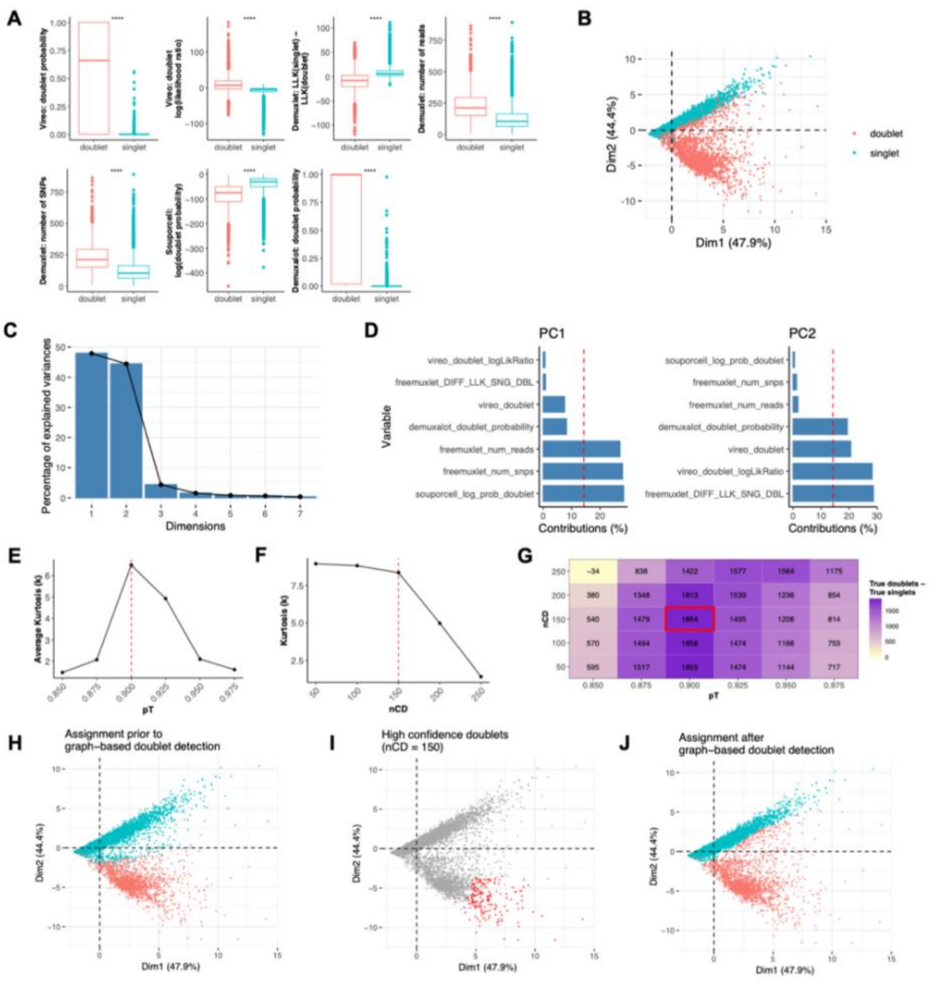


**Figure S5. Demonstration of the graph-based doublet detection component of the Ensemblex framework on an *in silico* pool with known ground-truth sample labels.** For this demonstration of the graph-based doublet detection component of the Ensemblex framework, we leveraged a computationally multiplexed pool comprising 24 induced pluripotent stem cell (iPSC) lines from genetically distinct individuals, 17,384 pooled cells, and a 15% doublet rate. **A)** Box plot showing the distribution of the doublet-related features across true singlets and true doublets. Wilcoxon rank-sum tests were used to compare the distribution of features across true singlets and doublets. **B)** Principal component analysis (PCA) using the doublet-related features showing true singlets and true doublets across the first two principal components (PCs). **C)** Scree plot showing the percentage of variance explained by the first seven PCs. **D)** Bar plots showing the percent contribution of each doublet-related variable to the first two PCs. **E)** Dot plot showing the average Pearson's measure of kurtosis (K) of the nearest neighbour frequency (fNN) density plots across percentile threshold (pT) values tested in the graph-based doublet detection parameter sweep. **F)** Dot plot showing K of the fNN density plots across varying numbers of confident doublet (nCD) tested in the parameter sweep for the optimal pT value (0.900). **G)** Heatmap showing the number of true doublets — the number of true singlets classified as doublets by the graph-based doublet detection component of the Ensemblex framework across various combinations of nCD and pT values tested in the parameter sweep. The red box shows the optimal nCD and pT values identified by the parameter sweep. **H)** PCA showing singlet and doublet classifications by Ensemblex after Step 1 (prior to graph-based doublet detection). **I)** PCA showing confident doublets identified by the graph-based doublet detection component of the Ensemblex framework. The optimal nCD are shown. **J)** PCA showing singlet and doublet classifications by Ensemblex after performing graph-based doublet detection. **** P-value < 0.0001.

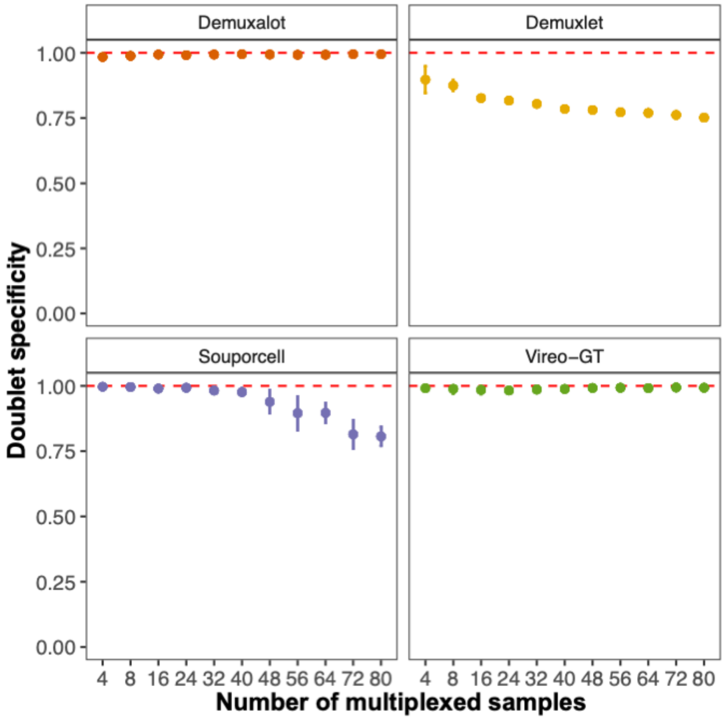


**Figure S6. Validating the ensemble-independent doublet detection component of the Ensemblex framework on *in silico* pools with known ground-truth sample labels.** The ensemble-independent doublet detection component of the Ensemblex framework retains the doublet labels from constituent demultiplexing tools that showed high doublet detection specificity on computationally multiplexed pools with known ground-truth sample labels. Namely, Ensemblex retains the doublet predictions made by Demuxalot and Vireo, by default. The doublet detection specificity of the individual tools was computed 96 *in silico* pools with known ground-truth sample labels ranging in size from 4 to 80 multiplexed induced pluripotent stem cell (iPSC) lines from genetically distinct individuals, averaging 17,396 cells per pool and a 15% doublet rate. The figure shows the mean doublet detection specificity of each constituent tool across replicates at a given sample size. Error bars represent one standard deviation from the mean.

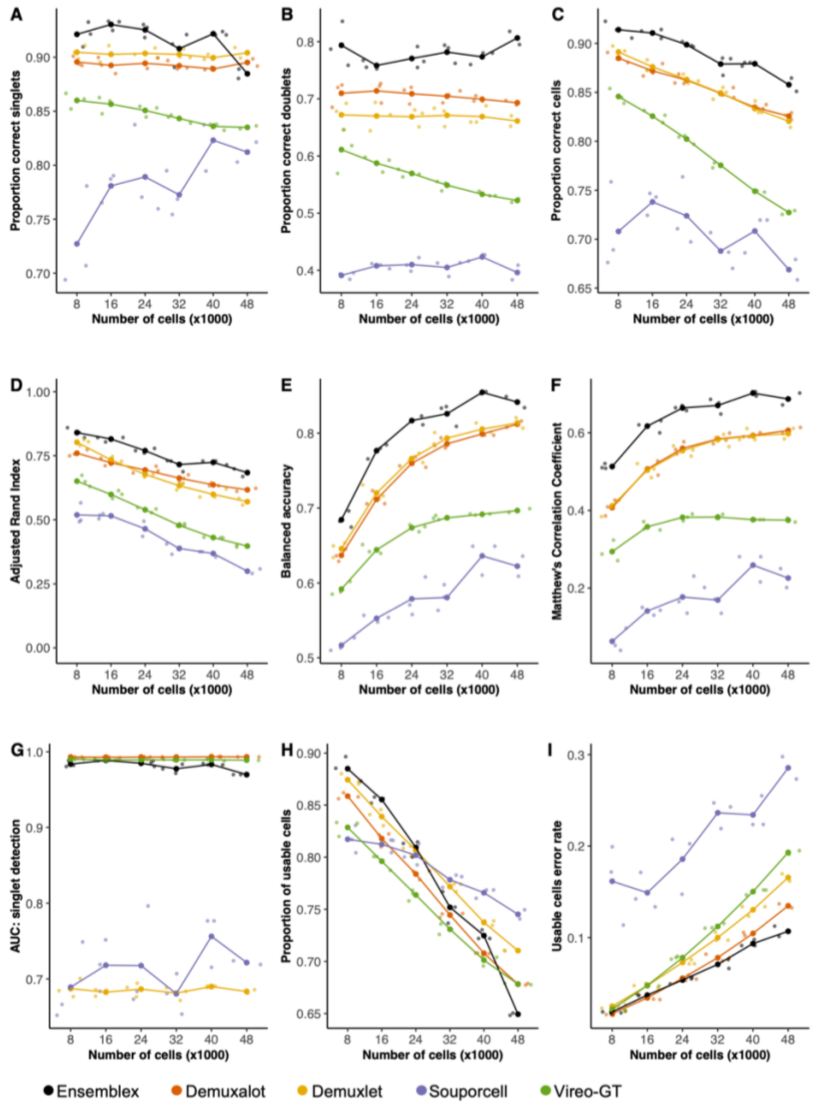


**Figure S7. Genetic demultiplexing performance with prior genotype information on *in silico* pools with varying numbers of pooled cells.** Genetic demultiplexing tools were evaluated on 18 *in silico* pools with known ground-truth sample labels ranging in size from 8,000 to 48,000 pooled cells with 24 multiplexed induced pluripotent stem cell (iPSC) lines from genetically distinct individuals and a doublet rate of 6% per 8,000 cells. A singlet was considered correctly classified if the assigned sample label matched the ground-truth sample label and the assignment probability exceeded the recommended threshold for the respective tool; a doublet was considered correctly classified if the assigned sample label matched the ground-truth sample label, regardless of the assignment probability. **A-I)** Line graphs showing the performance of Ensemblex and the individual genetic demultiplexing tools across evaluation metrics. The large dots show the mean value for each tool across replicates at a given pool size (n = 3 per pool size). **A)** Proportion of correctly classified singlets. **B)** Proportion of correctly classified doublets. **C)** Proportion of correctly classified cells. **D)** Adjusted Rand Index between each tool's sample labels and the ground- truth sample labels. **E)** Balanced accuracy of each tool. **F)** Matthew's Correlation Coefficient of each tool. **G)** Area under the receiver operating characteristic curve (AUC) of the singlet assignment probability for each tool. **H)** Proportion of usable cells returned by each tool. Usable cells were defined as singlets with an assignment probability exceeding the recommended threshold of the respective tool. **I)** Error rate amongst the usable cells returned by each tool; erroneous classifications comprised of true doublets labeled as singlets or true singlets assigned to the wrong sample.

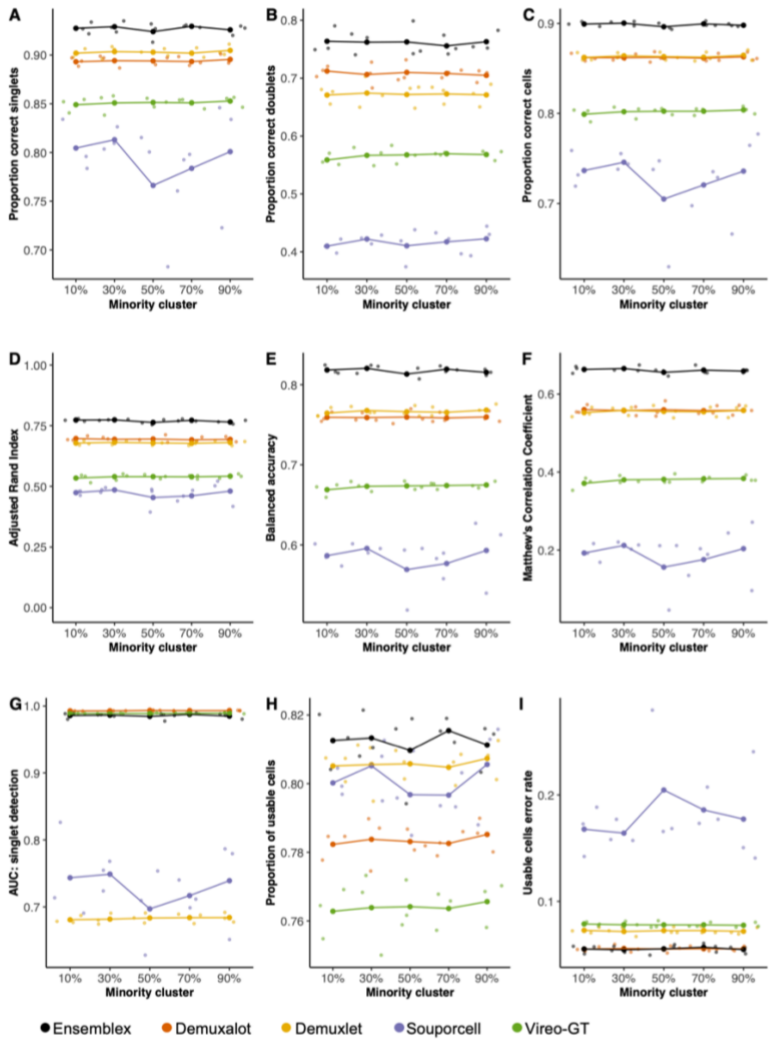


**Figure S8. Genetic demultiplexing performance with prior genotype information on *in silico* pools with an underrepresented cell line.** Genetic demultiplexing tools were evaluated on 15 *in silico* pools with known ground-truth sample labels, comprising 24 multiplexed induced pluripotent stem cell (iPSC) lines from genetically distinct individuals and a 15% doublet rate. For each pool, 23 iPSC lines contained 1,000 cells, while one randomly selected line showed various degrees of underrepresentation, namely 100 cells (10%), 300 cells (30%), 500 cells (50%), 700 cells (70%), or 900 cells (90%). A singlet was considered correctly classified if the assigned sample label matched the ground-truth sample label and the assignment probability exceeded the recommended threshold for the respective tool; a doublet was considered correctly classified if the assigned sample label matched the ground-truth sample label, regardless of the assignment probability. **A-I)** Line graphs showing the performance of Ensemblex and the individual genetic demultiplexing tools across evaluation metrics. The large dots show the mean value for each tool across replicates at a given degree of underrepresentation (n = 3 degree of underrepresentation). **A)** Proportion of correctly classified singlets. **B)** Proportion of correctly classified doublets. **C)** Proportion of correctly classified cells. **D)** Adjusted Rand Index between each tool's sample labels and the ground-truth sample labels. **E)** Balanced accuracy of each tool. **F)** Matthew's Correlation Coefficient of each tool. **G)** Area under the receiver operating characteristic curve (AUC) of the singlet assignment probability for each tool. **H)** Proportion of usable cells returned by each tool. Usable cells were defined as singlets with an assignment probability exceeding the recommended threshold of the respective tool. **I)** Error rate amongst the usable cells returned by each tool; erroneous classifications comprised of true doublets labeled as singlets or true singlets assigned to the wrong sample.

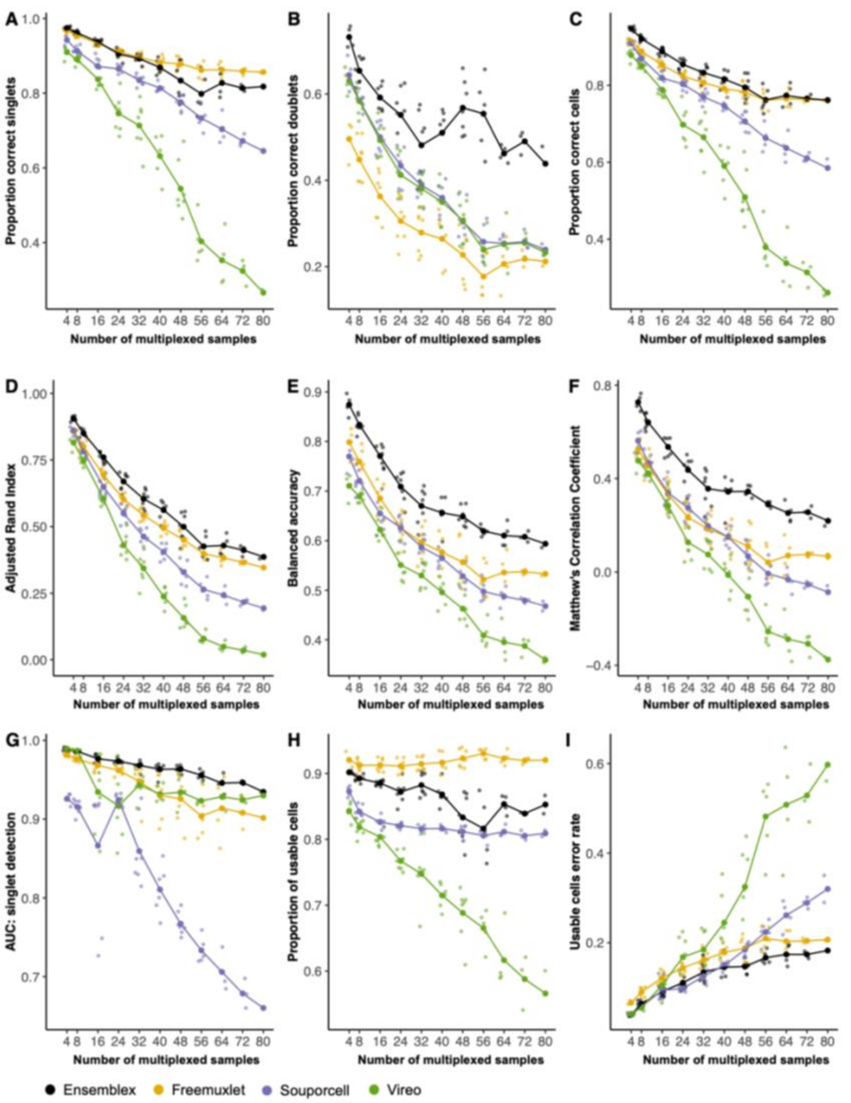


**Figure S9. Genetic demultiplexing performance without prior genotype information on *in silico* pools with varying numbers of multiplexed samples.** The genetic demultiplexing tools without prior genotype information were evaluated on 96 *in silico* pools with known ground-truth sample labels ranging in size from 4 to 80 multiplexed induced pluripotent stem cell (iPSC) lines from genetically distinct individuals, averaging 17,396 cells per pool and a 15% doublet rate. A singlet was considered correctly classified if the assigned sample label matched the ground-truth sample label and the assignment probability exceeded the recommended threshold for the respective tool; a doublet was considered correctly classified if the assigned sample label matched the ground-truth sample label, regardless of the assignment probability. **A-I)** Line graphs showing the performance of Ensemblex and the individual genetic demultiplexing tools across evaluation metrics. The large dots show the mean value for each tool across replicates at a given sample size (n = 9 per pool size). **A)** Proportion of correctly classified singlets. **B)** Proportion of correctly classified doublets. **C)** Proportion of correctly classified cells. **D)** Adjusted Rand Index between each tool's sample labels and the ground-truth sample labels. **E)** Balanced accuracy of each tool. **F)** Matthew's Correlation Coefficient of each tool. **G)** Area under the receiver operating characteristic curve (AUC) of the singlet assignment probability for each tool. **H)** Proportion of usable cells returned by each tool. Usable cells were defined as cells classified by singlets with an assignment probability exceeding the recommended threshold of the respective tool. **I)** Error rate amongst the usable cells returned by each tool; erroneous classifications comprised of true doublets labeled as singlets or true singlets assigned to the wrong sample.

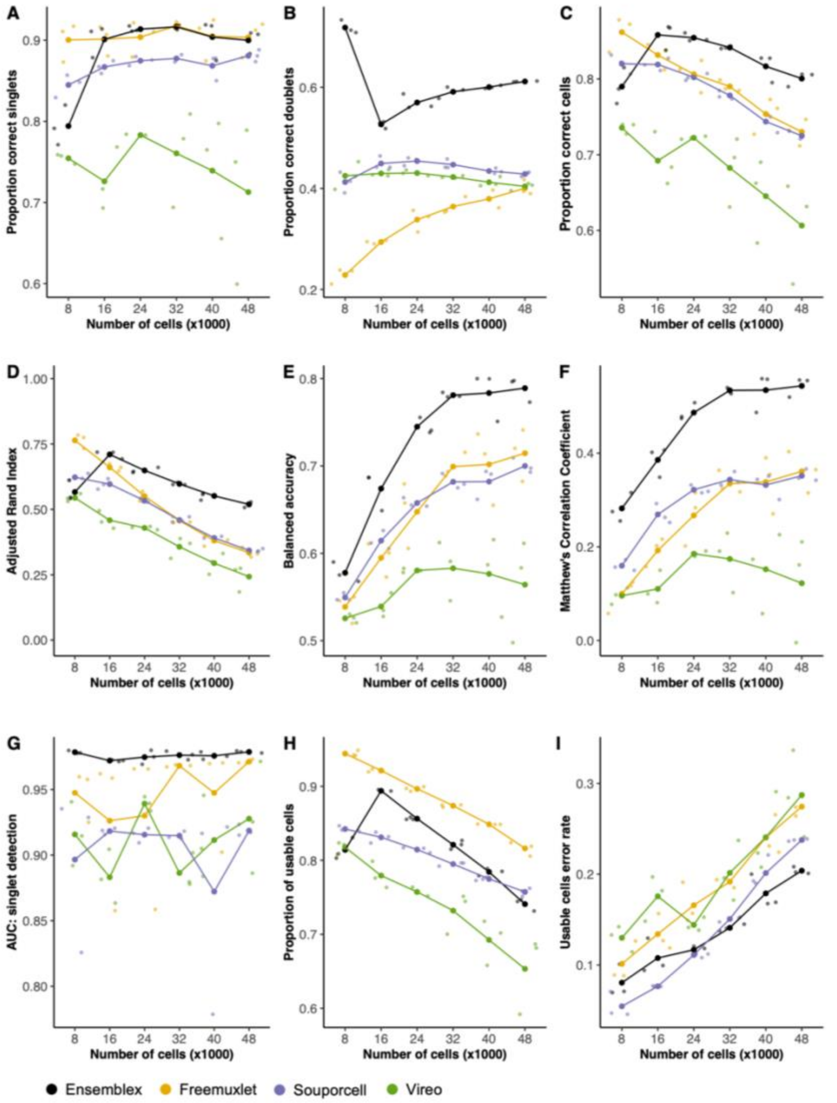


**Figure S10. Genetic demultiplexing performance without prior genotype information on *in silico* pools with varying numbers of pooled cells.** Genetic demultiplexing tools were evaluated on 18 *in silico* pools with known ground-truth sample labels ranging in size from 8,000 to 48,000 pooled cells with 24 multiplexed induced pluripotent stem cell (iPSC) lines from genetically distinct individuals and a doublet rate of 6% per 8,000 cells. A singlet was considered correctly classified if the assigned sample label matched the ground-truth sample label and the assignment probability exceeded the recommended threshold for the respective tool; a doublet was considered correctly classified if the assigned sample label matched the ground-truth sample label, regardless of the assignment probability. **A-I)** Line graphs showing the performance of Ensemblex and the individual genetic demultiplexing tools across evaluation metrics. The large dots show the mean value for each tool across replicates at a given pool size (n = 3 per pool size). **A)** Proportion of correctly classified singlets. **B)** Proportion of correctly classified doublets. **C)** Proportion of correctly classified cells. **D)** Adjusted Rand Index between each tool's sample labels and the ground- truth sample labels. **E)** Balanced accuracy of each tool. **F)** Matthew's Correlation Coefficient of each tool. **G)** Area under the receiver operating characteristic curve (AUC) of the singlet assignment probability for each tool. **H)** Proportion of usable cells returned by each tool. Usable cells were defined as singlets with an assignment probability exceeding the recommended threshold of the respective tool. **I)** Error rate amongst the usable cells returned by each tool; erroneous classifications comprised of true doublets labeled as singlets or true singlets assigned to the wrong sample.

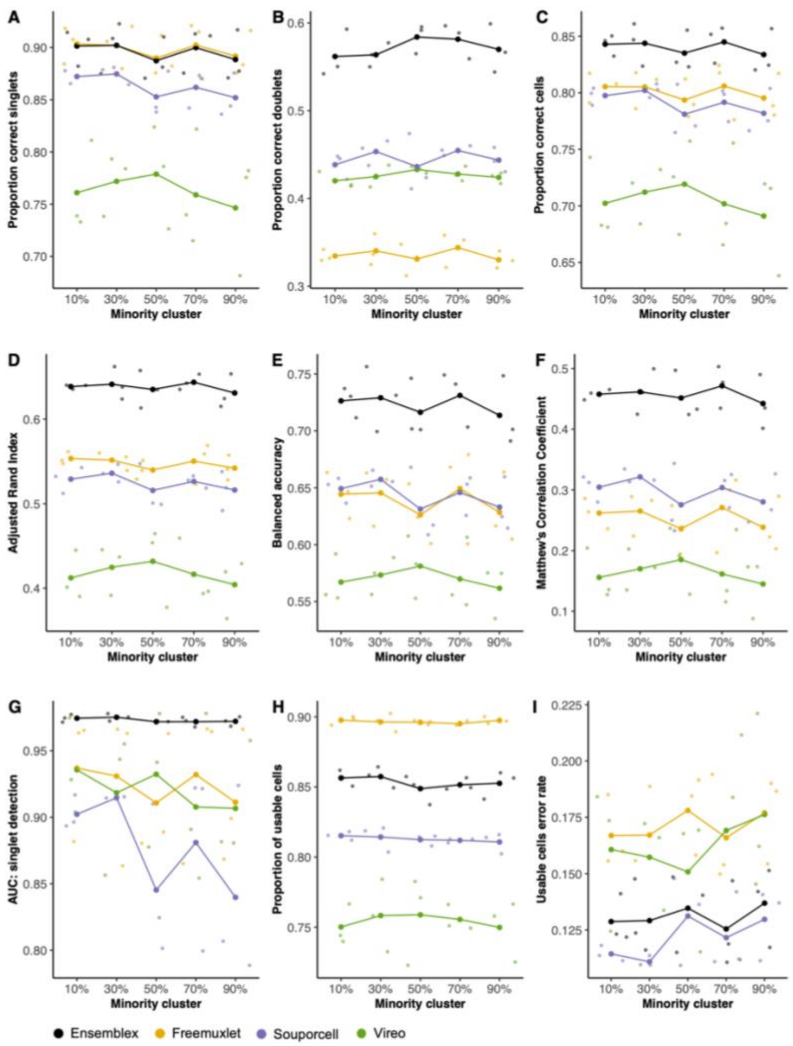


**Figure S11. Genetic demultiplexing performance without prior genotype information on *in silico* pools with an underrepresented cell line.** Genetic demultiplexing tools were evaluated on 15 *in silico* pools with known ground-truth sample labels, comprising 24 multiplexed induced pluripotent stem cell (iPSC) lines from genetically distinct individuals and a 15% doublet rate. For each pool, 23 iPSC lines contained 1,000 cells, while one randomly selected line showed various degrees of underrepresentation, namely 100 cells (10%), 300 cells (30%), 500 cells (50%), 700 cells (70%), or 900 cells (90%). A singlet was considered correctly classified if the assigned sample label matched the ground-truth sample label and the assignment probability exceeded the recommended threshold for the respective tool; a doublet was considered correctly classified if the assigned sample label matched the ground-truth sample label, regardless of the assignment probability. **A-I)** Line graphs showing the performance of Ensemblex and the individual genetic demultiplexing tools across evaluation metrics. The large dots show the mean value for each tool across replicates at a given degree of underrepresentation (n = 3 degree of underrepresentation). **A)** Proportion of correctly classified singlets. **B)** Proportion of correctly classified doublets. **C)** Proportion of correctly classified cells. **D)** Adjusted Rand Index between each tool's sample labels and the ground-truth sample labels. **E)** Balanced accuracy of each tool. **F)** Matthew's Correlation Coefficient of each tool. **G)** Area under the receiver operating characteristic curve (AUC) of the singlet assignment probability for each tool. **H)** Proportion of usable cells returned by each tool. Usable cells were defined as singlets with an assignment probability exceeding the recommended threshold of the respective tool. **I)** Error rate amongst the usable cells returned by each tool; erroneous classifications comprised of true doublets labeled as singlets or true singlets assigned to the wrong sample.

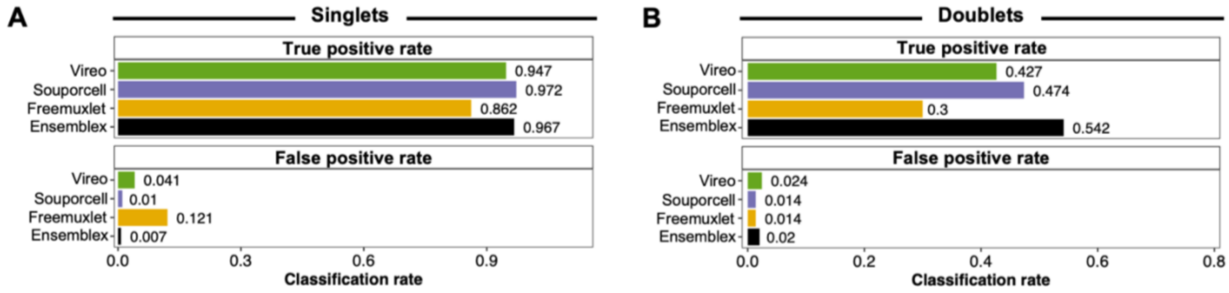


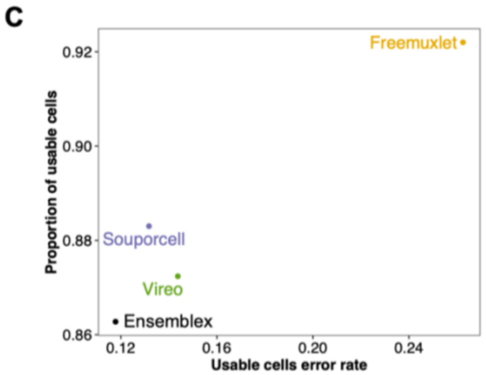


D

| | Ensemblex | Freemuxlet | Souporcell | Vireo |
|---|---|---|---|---|
| Adjusted Rand Index | 0.721 | 0.595 | 0.709 | 0.664 |
| Balanced Accuracy | 0.847 | 0.595 | 0.847 | 0.772 |
| Matthew's Correlation Coefficient | 0.593 | 0.175 | 0.556 | 0.451 |
| AUC singlet detection | 0.924 | 0.600 | 0.587 | 0.928 |

**Figure S12. Evaluating Ensemblex without prior genotype information on experimentally multiplexed cells using donor-specific oligonucleotide labels as a proxy for ground-truth.** Non-small cell lung cancer (NSCLC) dissociated tumor cells from seven individuals were pooled and labelled with donor-specific oligonucleotide-labels. Cells were demultiplexed according to their expression of donor-specific oligonucleotide labels by HTOdemux; HTOdemux's sample labels were used as a proxy for ground truth. True positives (TP) singlets were defined as cells classified as singlets by both HTOdemux and Ensemblex with matching sample labels; false positives (FP) singlets were defined as cells classified as singlets by both HTOdemux and Ensemblex but assigned to different donors. TP doublets were defined as cells classified as doublets by both HTOdemux and Ensemblex; FP doublets were defined as cells classified as singlets by HTOdemux and doublets by Ensemblex; false negatives (FN) doublets were defined as cells classified as doublets by HTOdemux and singlets by Ensemblex. **A)** T-distributed Stochastic Neighbor Embedding (t-SNE) visualization of HTOdemux's sample labels. **B)** T-SNE visualization of Ensemblex's demultiplexing performance using HTOdemux's sample labels as ground truth for singlets (left) and doublets (right). **C)** Bar plots showing the singlet TP and FP rates for each genetic demultiplexing tool using HTOdemux's sample labels as ground truth. **D)** Bar plots showing the doublet TP and FP rates for each genetic demultiplexing tool using HTOdemux's sample labels as ground truth. **E)** Scatter plot showing the proportion of usable cells (confidently classified singlets) and the corresponding usable cell error rate for each genetic demultiplexing tool. **F)** Adjusted Rand Index, balanced accuracy, Matthew's Correlation Coefficient, and area under the receiver operating characteristic curve (AUC) of the singlet assignment probability for each genetic demultiplexing tool.

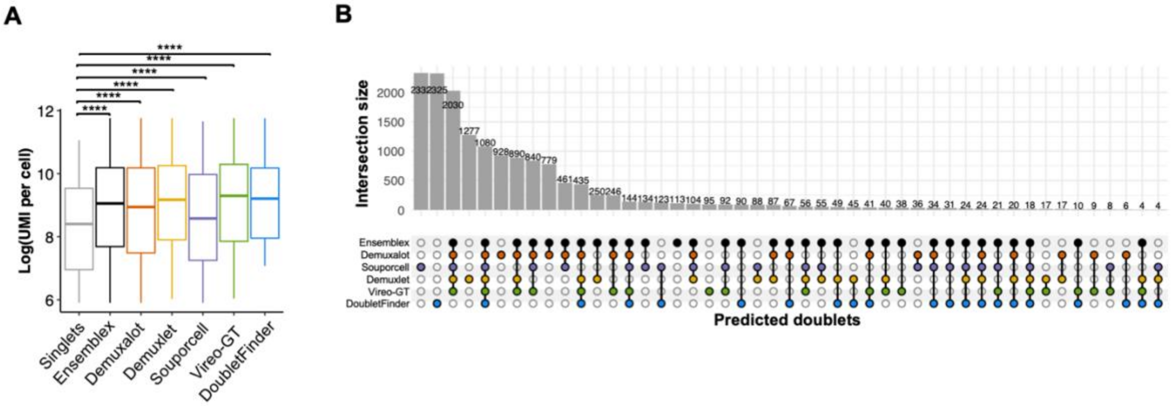


**Figure S13. Doublets identified by individual tools across the dopaminergic neuron dataset. A)** Boxplot showing the **distribution** of unique molecular identifiers (UMI) per cell across doublets identified by each tool and consensus singlets — cells ubiquitously assigned as singlets by each tool. Wilcoxon-rank sum tests compared the distribution of UMI per cell across singlets to that of the doublets identified by each tool. **B)** Upset plot showing the intersection of doublets identified by each tool. **** Adjusted P-value < 0.0001.

**Table S1. Evaluating the contribution of each component of the Ensemblex framework to the overall demultiplexing accuracy with prior genotype information on experimentally multiplexed cells using donor-specific oligonucleotide labels as a proxy for ground-truth.**

| | Ensemblex demultiplexing performance | | |
|---|---|---|---|
| | After Accuracy-weighted probabilistic ensemble | After Graph-based doublet detection | After Ensemble-independent doublet detection |
| Singlet true positive rate | 0.974 | 0.973 | 0.969 |
| Doublet true positive rate | 0.518 | 0.589 | 0.662 |

Evaluation of Ensemblex was based on HTOdemux's sample labels. Singlet true positive rate: proportion of singlets identified by HTOdemux that had the same sample labels as Ensemblex; doublet true positive rate: proportion of doublets identified by HTOdemux that were identified as doublets by Ensemblex.

**Table S2. Evaluating the contribution of each component of the Ensemblex framework to the overall demultiplexing accuracy without prior genotype information on experimentally multiplexed cells using donor-specific oligonucleotide labels as a proxy for ground-truth.**

| | Ensemblex demultiplexing performance | | |
|---|---|---|---|
| | After Accuracy-weighted probabilistic ensemble | After Graph-based doublet detection | After Ensemble-independent doublet detection |
| Singlet true positive rate | 0.972 | 0.971 | 0.967 |
| Doublet true positive rate | 0.417 | 0.485 | 0.542 |

Evaluation of Ensemblex was based on HTOdemux's sample labels. Singlet true positive rate: proportion of singlets identified by HTOdemux that had the same sample labels as Ensemblex; doublet true positive rate: proportion of doublets identified by HTOdemux that were identified as doublets by Ensemblex.

**Table S3. Description of technical replicates from the dopaminergic neuron dataset.**

| Sequencing timepoint | Technical replicate | *n* multiplexed samples | *n* cells | ENA accession IDs |
|---|---|---|---|---|
| Day 11 | 1 | 22 | 9,314 | ERR4700135<br>ERR4700136<br>ERR4700137<br>ERR4700138 |
| Day 11 | 2 | 22 | 8,291 | ERR4700139<br>ERR4700140<br>ERR4700141<br>ERR4700142 |
| Day 11 | 3 | 22 | 10,171 | ERR4700143<br>ERR4700144<br>ERR4700145<br>ERR4700146 |
| Day 30 | 1 | 22 | 10,297 | ERR4700183<br>ERR4700184<br>ERR4700185<br>ERR4700186 |
| Day 30 | 2 | 22 | 9,486 | ERR4700187<br>ERR4700188<br>ERR4700189<br>ERR4700190 |
| Day 30 | 3 | 22 | 10,590 | ERR4700191<br>ERR4700192<br>ERR4700193<br>ERR4700194 |
| Day 52 | 1 | 22 | 8,712 | ERR4700199<br>ERR4700200<br>ERR4700201<br>ERR4700202 |
| Day 52 | 2 | 22 | 8,645 | ERR4700203<br>ERR4700204<br>ERR4700205<br>ERR4700206 |
| Day 52 | 3 | 22 | 9,240 | ERR4700211<br>ERR4700212<br>ERR4700213<br>ERR4700214 |

Pooled cultures of induced pluripotent stem cell (iPSC) lines from 22 healthy donors were differentiated towards a dopaminergic neuron (DaN) fate and sequenced on days 11, 30, and 52 of differentiation by Jerber et al. For the analysis we used three technical replicates for each sequencing timepoint. ENA: European Nucleotide Archive.

**Table S4. Demographic and clinical information for the subjects analyzed in the neural stem cell dataset.**

| Cell line | Clones | Cell type of origin | Diagnosis | Age | Biological sex |
|---|---|---|---|---|---|
| K001 | i6 & i9 | Keratinocytes | Healthy control | 15 | Male |
| K005 | z12 & z13 | Keratinocytes | Healthy control | 16 | Male |
| K011 | c6 & c10 | Keratinocytes | Healthy control | 16 | Male |
| K013 | c20 & c20 | PBMCs | Healthy control | 9 | Female |
| K015 | c1 & c9 | PBMCs | Healthy control | 13 | Male |
| 205 | c2 | Lymphoblastoids | ADHD | 16 | Male |
| MR001 | x3 & x15 | Keratinocytes | ADHD | 15 | Male |
| MR010 | c3 & c18 | Keratinocytes | ADHD | 9 | Male |
| MR012 | c11 | PBMCs | ADHD | | Male |
| MR013 | c3 & c13 | PBMCs | ADHD | 16 | Male |
| MR014 | c12 & c27 | PBMCs | ADHD | 13 | Male |
| MR030 | c1 & c2 | PBMCs | ADHD | 9 | Female |
| NR002 | c2 & c21 | PBMCs | ADHD | 13 | Male |
| 308 | c1 | Lymphoblastoids | ADHD | 15 | Female |

**Table S5. Neural stem cell samples submitted for single-cell RNA sequencing and the cell lines composing their respective pools.**

| Experiment | Cell type | Pool | Cell line |
|---|---|---|---|
| 1 | NSCs | 1.1 | K015 c1 |
| | | | K011 c10 |
| | | | K013 c20 |
| | | | NR002 c21 |
| | | | MR010 c3 |
| | | | MR014 c27 |
| | | | MR030 c1 |
| | | 1.2 | K001 i6 |
| | | | K005 z12 |
| | | | K008 i13 |
| | | | MR023 c17 |
| | | | MR013 c13 |
| | | | MR001 x3 |
| 2 | NSCs | 2.1 | NR002 c2 |
| | | | K013 c39 |
| | | | MR014 c12 |
| | | | K011 c6 |
| | | | K015 c9 |
| | | | K008 i44 |
| | | | 205 c2 |
| | | | 308 c1 |
| | | 2.2 | MR012 c11 |
| | | | MR010 c18 |
| | | | MR001 x15 |
| | | | MR030 c2 |
| | | | MR023 c22 |
| | | | K001 i9 |
| | | | K005 z13 |
| | | | MR013 c3 |

**Table S6. Distribution of putative cell types across individuals with ADHD and controls from the neural stem cell dataset according to assignments by the genetic demultiplexing tools.**

| | **Ensemblex** | | **Demuxalot** | | **Demuxlet** | | **Souporcell** | | **Vireo-GT** | |
|---|---|---|---|---|---|---|---|---|---|---|
| | ADHD | Ctrl | ADHD | Ctrl | ADHD | Ctrl | ADHD | Ctrl | ADHD | Ctrl |
| Glia | 274 | 2355 | 265 | 2304 | 249 | 2049 | 514 | 3266 | 339 | 2521 |
| NSC-SOX2 | 579 | 4798 | 590 | 4812 | 582 | 4120 | 864 | 4571 | 678 | 4497 |
| NPC-POU5F1 | 313 | 3874 | 279 | 3253 | 295 | 3035 | 595 | 5574 | 283 | 2498 |
| Neuron-DCX | 680 | 3755 | 630 | 3740 | 657 | 3259 | 937 | 3274 | 745 | 3039 |
| Neuron-GRIA1 | 413 | 1974 | 328 | 1647 | 349 | 1657 | 621 | 1654 | 207 | 675 |
| NPC-S100B | 385 | 2122 | 378 | 2117 | 370 | 1964 | 563 | 2027 | 447 | 1748 |
| NPC-MEF2C | 65 | 693 | 63 | 662 | 66 | 626 | 161 | 1022 | 84 | 661 |
| NSC-APOA1 | 30 | 309 | 25 | 322 | 32 | 263 | 46 | 309 | 35 | 316 |
